# INCORPORATING A DYNAMIC GENE-BASED PROCESS MODULE INTO A CROP SIMULATION MODEL

**DOI:** 10.1101/2021.02.02.429409

**Authors:** Fabio A.A. Oliveira, James W. Jones, Willingthon Pavan, Mehul Bhakta, C. Eduardo Vallejos, Melanie J. Correll, Kenneth J. Boote, José M.C. Fernandes, Carlos A. Hölbig, Gerrit Hoogenboom

## Abstract

Dynamic crop simulation models are tools that predict plant phenotype grown in specific environments for genotypes using genotype-specific parameters (GSPs), often referred to as “genetic coefficients.” These GSPs are estimated using phenotypic observations and may not represent “true” genetic information. Instead, estimating GSPs requires experiments to measure phenotypic responses when new cultivars are released. The goal of this study was to evaluate a new approach that incorporates a dynamic gene-based module for simulating time-to-flowering for common bean (*Phaseolus vulgaris* L.) into an existing dynamic crop model. A multi-environment study conducted in 2011 and 2012 included 187 recombinant inbred lines (RILs) from a bi-parental bean family to measure the effects of quantitative trait loci (QTL), environment (E), and QTL×E interactions across five sites. The dynamic mixed linear model from Vallejos et al. (2020) was modified in this study to create a dynamic module that was then integrated into the CSM-CROPGRO-Drybean model. This new hybrid crop model, with the gene-based flowering module replacing the original flowering component, requires allelic makeup of each genotype being simulated and daily E data. The hybrid model was compared to the original CSM model using the same E data and previously estimated GSPs to simulate time-to-flower. The integrated gene-based module simulated days of first flower agreed closely with observed values (root mean square error of 2.73 days and model efficiency of 0.90) across the five locations and 187 genotypes. The hybrid model with its gene-based module also described most of the G, E and G×E effects on time-to-flower and was able to predict final yield and other outputs simulated by the original CSM. These results provide the first evidence that dynamic crop simulation models can be transformed into gene-based models by replacing an existing process module with a gene-based module for simulating the same process.

## 1. Introduction

Scientific advances in understanding plant genes combined with advances in technologies for rapidly and inexpensively identifying genetic makeup of plants (Thomson 2014; Rasheed *et al*. 2017) have fueled considerable interest in using genetic information to predict plant phenotypes. Analytical tools are now available to identify the genes that are associated with the variation in different plant traits. These bioinformatics tools also can identify important gene-by-environment (G × E) interactions that contribute to observed variations in specific traits (Yin *et al*. 2018). Rapid progress in genome-wide association studies (GWAS) has enabled researchers to identify genes associated with variations in human diseases (Bush and Moore 2012).

Genome-wide prediction models that use GWAS also have become powerful tools for improving crops such as tropical rice (e.g., Spindel *et al*. 2016). The GWAS approach has been implemented in recent work in other crops (Brown *et al*. 2014; Huang and Han 2014; Cooper *et al*. 2016). Scientists use statistical methods, such as single locus analysis based on ANOVA, linear regression, and mixed linear regression models, to detect a gene or gene combinations associated with variations in a phenotypic trait (White and Hoogenboom 2003; White 2006; Yin and Struik 2010). These tools also assist geneticists and plant breeders for prediction and selection of lines to improve crop yield.

Concepts have been under development since the early 1970s for predicting crop yield variations using dynamic models as affected by environmental conditions and management scenarios, and to some variations among cultivars (Jones *et al*., 2016; Thorburn et al., 2018). Differences among cultivars are represented by empirical genotype-specific parameters (GSPs). However, these models do not use information on variations in genes among the cultivars. Instead, the GSPs for each genotype must be estimated using data from laboratory or field studies (Hunt *et al*. 1993; Anothai *et al*. 2008; Buddhaboon *et al*. 2018).

Recognizing the potential for introducing genetic information into crop models, White and Hoogenboom (1996, 2003) showed that some of the BEANGRO model’s GSPs (Hoogenboom *et al*. 1992, 1994) could be estimated as linear functions of genetic information. This approach was also used by Messina et al. (2006) for the CROPGRO-Soybean model and by other researchers for different crops (Yin *et al*. 2000, 2003; Reymond *et al*. 2003; Hammer *et al*. 2010; Gu *et al*. 2014). Furthermore, this approach of relating existing crop model GSPs to molecular markers was shown to provide better yield predictions than that of a statistical model for maize (Technow *et al*. 2015). More recently, Wallach *et al*. (2018) showed that genetic effects on rate of progress to first flower in common bean can be estimated using field data from a multi-environmental trial containing a large number of genotypes.

Although GSPs can provide high levels of prediction in crop models when they are independently estimated for each genotype, these parameter may not accurately represent the genetic architecture of the associated crop phenotype or process (Hwang *et al*. 2017). Acharya *et al*. (2017) found that commonly-used approaches for estimating GSPs for the Cropping System Model (CSM)-CROPGRO-Drybean model (Boote *et al*. 1998; Jones *et al*. 2003; Hoogenboom *et al*. 2019a) resulted in considerable equifinality among estimated GSPs, which means that multiple sets of possible GSP values produced very similar responses. This was demonstrated by Acharya et al. (2017) using a synthetic population based on known GSPs that were used to generate synthetic field data. Then, blind estimates of GSPs using those synthetic data differed from original values. Even though new GSPs reliably predicted crop growth and yield, the procedure was unable to recover the original GSP values.

The previously discussed studies contain relationships and assumptions made by the original crop model developers, including the functional forms used to describe the E and G effects on predicted dynamic rates. As a result, this use of existing relationships makes it difficult to identify G and G × E effects from field data, which can be seen in the expanded original model form published by Wallach *et al*. (2018). Note that this expanded functional form inherently includes many G × E interaction terms that may or may not exist. Incorporating genetic information into an original model’s functional form entangles G, E, and G × E effects and, thus, does not enable one to study interactive G × E effects on the rate of progress toward flowering. Furthermore, we have learned that there are variations among genotypes that were not captured in the original model formulations and associated assumptions (Boote *et al*. 2003; Boote *et al*. 2013; Acharya *et al*. 2017). One example is that some combinations of genes may result in different responses to temperature than others, whereas the assumptions imbedded in the existing models mostly assume that temperature responses of all genotypes are the same.

Another issue is that the gene-based approach that thus far has evolved in the crop modeling community has not been widely embraced by the genetics community, nor have the analytical approaches used by geneticists to predict genetic effects on crop traits been adopted by the crop modeling community. There have been limited interactions between these science communities that might lead to more rapid advances in gene-based modeling. Hwang *et al*. (2017) concluded that comprehensive gene-based crop models may be developed using existing crop models by replacing existing component dynamic modules with gene-based modules as they are developed. Recent progress has helped identify a possible pathway to help converge these communities. A multi-environment trial (MET) that was conducted in 2011 and 2012 included 187 common bean (*Phaseolus vulgaris* L.) recombinant inbred lines (RIL) from a bi-parental family. As part of this study, significant QTLs controlling the time to flowering in the RIL population were identified (Bhakta *et al*. 2015, 2017), which provided an opportunity to model QTL and environmental effects on the time to flowering. This study included geneticists, biostatisticians, and crop modelers asking questions about which genes affected different growth and development processes across environments and what G × E interactions were important. One of the outcomes of this study was a QTL-based mixed model to determine the G and G × E interactions in order to build a predictive model for the time-to-flowering trait (Bhakta *et al*. 2017).

Vallejos *et al*. (2020) described how one can use a statistical model to develop a dynamic mixed model that can predict the time to first flower phenotype based on a daily development rates. They developed a model that predicts the daily development rate and discussed its potential integration into an existing dynamic crop model that responds to varying environmental conditions. However, integration of this model was not attempted by Vallejos *et al*. (2020). There could be unknown or implicit assumptions in the original crop model that might lead to erratic responses that would have to be identified and addressed when a gene-based module is integrated into an existing dynamic crop model that does not rely on genetic data inputs.

The goal of this study was, therefore, to address the questions of how this type of integration can be done in a comprehensive dynamic crop model and what complications and limitations are likely to occur. The first objective was to develop and integrate a dynamic statistical gene-based module into the CSM-CROPGRO-Drybean model to predict the time of first flower appearance using data obtained from the MET bean studies. The second objective was to demonstrate potential applications of the hybrid dynamic model as a breeding tool for studying the G × E interactions using sensitivity analysis and for simulating yield.

## 2. Materials & Methods

### 2.1. Genotype Population

The bean MET was conducted to collect the time-to-flowering phenotypes of a RIL population from a cross between the Andean bean cultivar, Calima, and a Mesoamerican cultivar, Jamapa (Bhakta *et al*. 2017). The Calima parent is a large-seeded, mottled bean Colombian cultivar with a determinate growth habit, while Jamapa is a small, black-seeded Mexican cultivar with an indeterminate growth habit. The RIL population was developed through single seed descent for 10 generations, followed by bulk propagation for an additional three generations (F_11:14_) giving rise to 187 RILs. Further details for this RIL population can be found in Bhakta *et al*. (2017), while the QTL-based linkage is described by Bhakta *et al*. (2015).

### 2.2. Experimental sites

We used the data from the MET study that included five locations, 187 RILs, and two parents reported by Bhakta *et al*. (2017). The five sites had been selected to provide contrasting environmental growing conditions, especially those related to temperature and photoperiod. Three of the five sites are located in the USA: Prosper, North Dakota (ND); Citra, Florida (FL); and Isabela, Puerto Rico (PR), while the other two sites are located in Colombia: Palmira, (PA), and Popayan, (PO). Fig. 1 and Table 1 summarize the seasonal temperature, day length, and solar radiation for the five sites in the MET study, which are the main environmental variables that affect the time to flowering in common bean. Prosper (ND) has longer days than the other environments, while Palmira and Popayan are close to the equator and have short days. Within Colombia, Popayan, the coolest site, is located at an elevation of 1,800 m, while Palmira, the warmer site, is located at a 1,000 m elevation.

**Figure 1.**
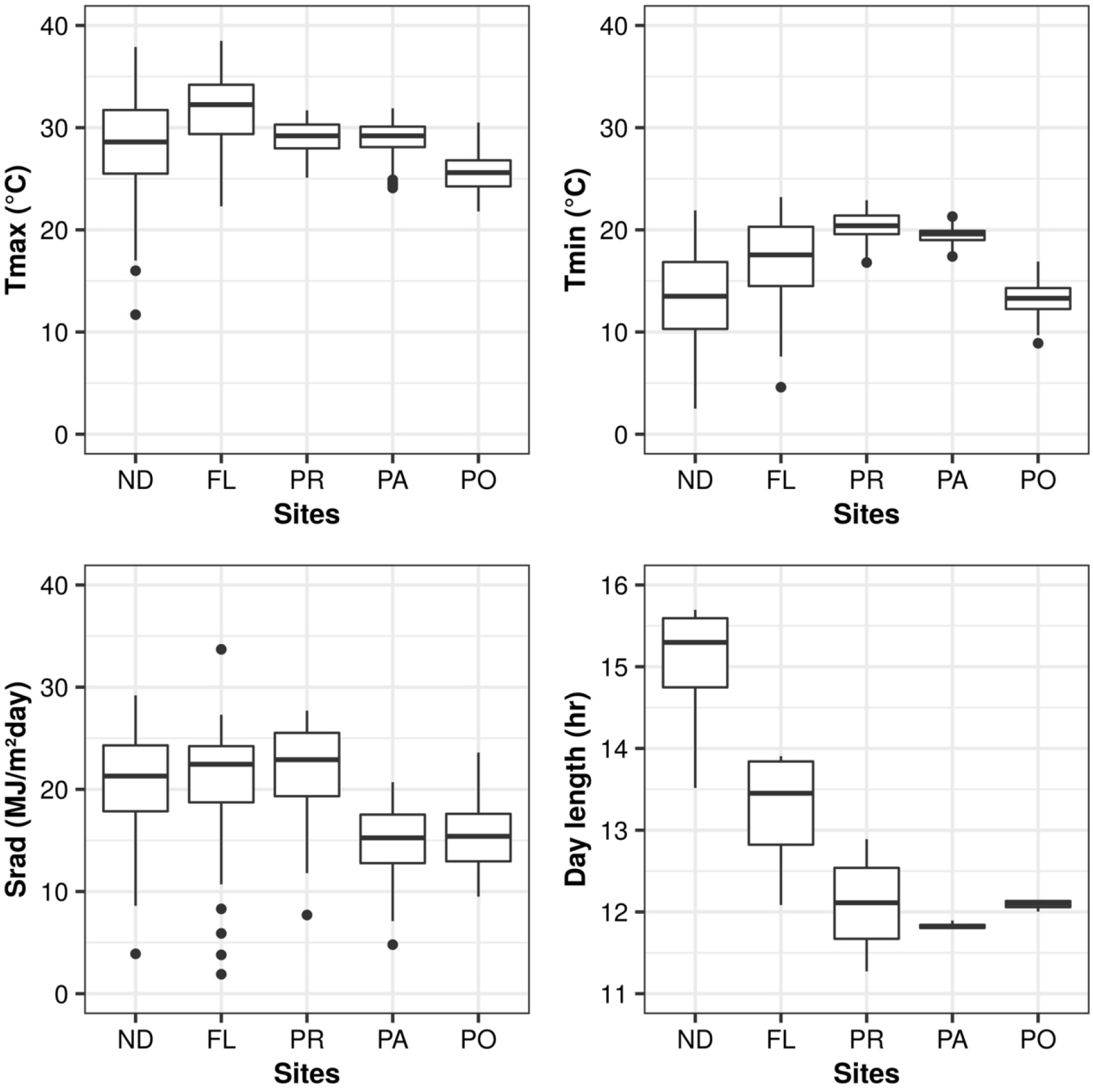
Boxplots of environmental variables observed for all five sites: Prosper, North Dakota (ND); Citra, Florida (FL); Isabella, Puerto Rico (PR); Palmira, Colombia (PA); Popayan, Colombia (PO). The boxplots show the distribution of daily values of maximum temperature (top left), daily minimum temperature (top right), daily solar radiation (bottom left) and day length (bottom right). The day length for all locations is based on the calculations of the CSM-CROPGRO-Drybean model.

**Table 1.** Summary of the meteorological and geographical data for each site in the multi-environment trial as reported by Bhakta *et al*. (2017), except for the computed day length that is based on the Cropping System Model (CSM).

|  | **ND** | **FL** | **PR** | **PA** | **PO** |
| --- | --- | --- | --- | --- | --- |
| **Location** | Prosper,<br>North<br>Dakota, USA | Citra, Florida,<br>USA | Isabella,<br>Puerto Rico,<br>USA | Palmira,<br>Colombia | Popayan,<br>Colombia |
| **Latitude /** | 47° 00' N / | 29° 39' N / | 18° 28' N / | 03° 29' N / | 02° 25' N / |
| **Longitude** | 96° 47' W | 82° 06' W | 61° 02' W | 76° 81' W | 76° 62' W |
| **Elevation (m)** | 280 | 31 | 128 | 1000 | 1800 |
| **Earliest first flowering date** | Jun-30-2012 | Apr-26-2012 | Mar-6-2012 | Dec-9-2011 | Apr-28-2012 |
| **Latest first flowering date** | Jul-26-2012 | May-16-2011 | Mar-22-2012 | Dec-24-2011 | May-16-2012 |
| **Seasonal Maximum** | 27 | 32 | 29 | 28 | 25 |
| **Temperature (°C)<sup>a</sup>** |  |  |  |  |  |
| **Seasonal Minimum** | 13 | 18 | 19 | 19 | 13 |
| **Temperature (°C)<sup>a</sup>** |  |  |  |  |  |
| **Day-Length (hh:mm)<sup>b</sup>** | 15:29 | 12:41 | 11:33 | 11:49 | 12:03 |
| **Solar Radiation (MJ m<sup>-2</sup> d<sup>-1</sup>)<sup>c</sup>** | 21.0 | 20.6 | 21.5 | 13.8 | 15.0 |

<sup>a</sup>Growing season average values for maximum and minimum temperature.
<sup>b</sup>Average day length data from sowing to first flower as computed by the CSM of the Decision Support System for Agrotechnology Transfer (DSSAT).
<sup>c</sup>Growing season average daily total solar radiation.

The experiment was conducted in 2011 and 2012, depending on the site. Each RIL and the two parents were grown in three replicated plots per site, with between 35 and 50 plants per plot. Six individual plants per plot were tagged at the V1 (first trifoliate opening) stage to record the vegetative and reproductive growth stages, resulting in 18 observations per genotype per site for each observation day. The plants were monitored daily to determine the date for each individual plant when first flowering occurred.

### 2.3. Dynamic mixed linear model

Vallejos *et al*. (2020) described procedures used to develop a dynamic mixed linear model to determine the rate of progress towards first flowering. This model was based on earlier work that was conducted by Bhakta *et al*. (2017) who fitted a statistical mixed linear model to predict time-to-flowering of the RILs based on QTL information and the mean environmental variables for each of the five sites. The Bhakta *et al*. (2017) model used a linear function for the effects of maximum and minimum temperature, day length, and solar radiation, each averaged over the duration between sowing and first flowering, twelve QTLs, five QTL × E factors, and one QTL × QTL factor. This non-dynamic model was able to describe 89% of the observed variability among the five locations and 187 RILs, with a root mean square error (RMSE) of 2.52 days.

Vallejos *et al*. (2020) used a similar approach to that used by Bhakta *et al*. (2017) to develop their dynamic model (Supporting Information – Fig. S1). First, the time to first flower data for all RILs and environment combinations were transformed into a development rate toward first flower appearance, calculated as rate = 1/(days to first flower). This approach requires the implementation of a function that predicts the daily development rate towards the time to first flower.

We designed a new module (DMLM; see Table 2 for abbreviations) by converting the Vallejos et al. (2020) model into a form that could be integrated with the original CSM-CROPGRO-Drybean model. The dynamic module computes the fraction of daily progress towards flowering based on the developmental rate that is controlled by genotype and daily environmental conditions. The time-to-flowering is determined when the cumulative addition of the daily progress time steps reaches unity. Eq. (1) shows the DMLM module that contains four environmental variables, one QTL × QTL interaction and seven QTL × E interactions.

**Table 2.**
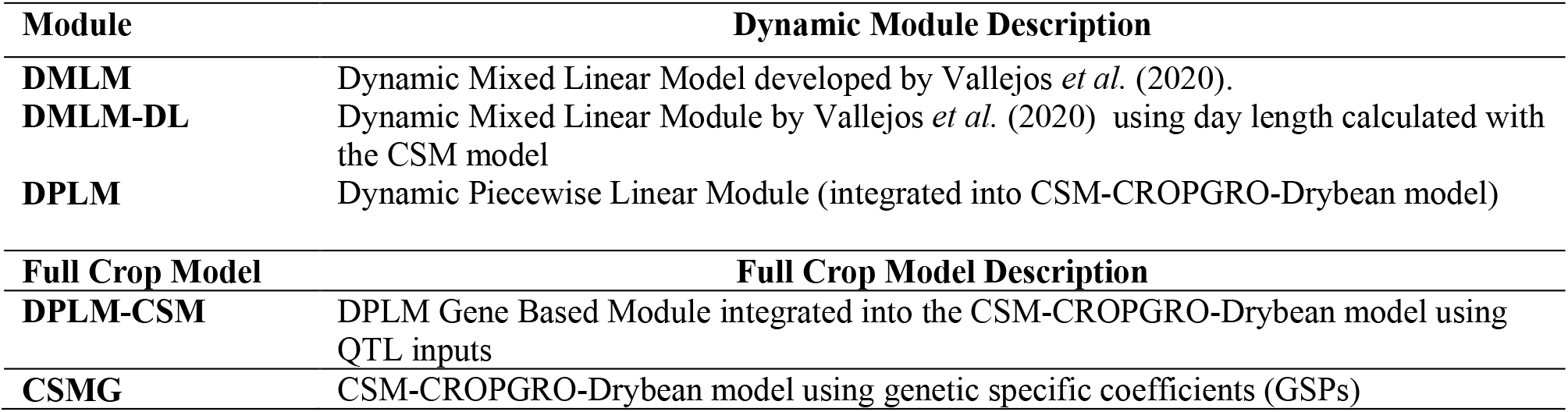
Description of model abbreviations.

| **Module** | **Dynamic Module Description** |
| --- | --- |
| **DMLM** | Dynamic Mixed Linear Model developed by Vallejos *et al.* (2020). |
| **DMLM-DL** | Dynamic Mixed Linear Module by Vallejos *et al.* (2020) using day length calculated with the CSM model |
| **DPLM** | Dynamic Piecewise Linear Module (integrated into CSM-CROPGRO-Drybean model) |
| **Full Crop Model** | **Full Crop Model Description** |
| **DPLM-CSM** | DPLM Gene Based Module integrated into the CSM-CROPGRO-Drybean model using QTL inputs |
| **CSMG** | CSM-CROPGRO-Drybean model using genetic specific coefficients (GSPs) |

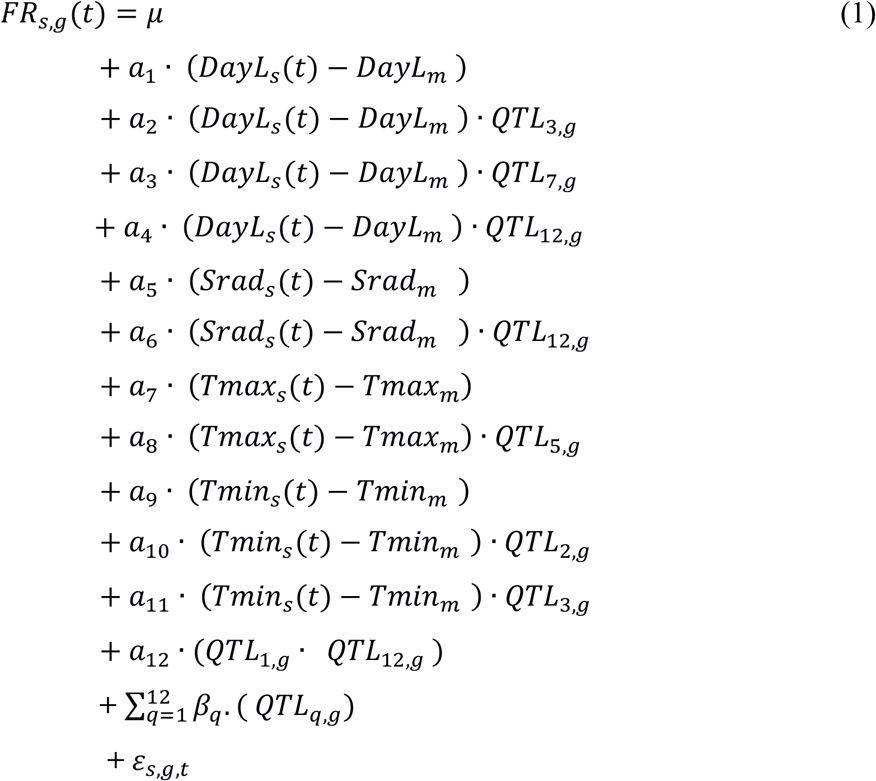

Where *FR*_*s,g*_(t) is the rate of progress to flowering (1/d) for the *g*^th^ genotype for the *s*^th^ site at time *t* (in days). *μ* represents the overall mean value of the daily development rates across all RILs and sites in the MET dataset. In the linear function (Eq. 1), RILs were treated as random effects and all remaining factors were considered as fixed effects. The variance-covariance structure that was used was unstructured, which is the default in the lme4 R-package. Note that Eq. (1) uses E variables that are centered on the mean values from the MET study for each RIL, based on the Vallejos *et al*. (2020) model. The first terms express the effects of the four environmental variables and the QTL-by-E effects, in which the variables *a*_*1*_ through *a*_*11*_ are estimated coefficients that quantify those effects, *DayL*_*s*_*(t)* is day length on each day *t* of the experiment at site *s*. Similarly, *Srad*_*s*_*(t)* is daily solar radiation (MJ/m^2^), *Tmax*_*s*_*(t)* is daily maximum temperature (°C), and *Tmin*_*s*_*(t)* is daily minimum temperature (°C) for each day *t* at site *s*. Mean values for each environmental variable (Eq. 1) were used as constants to center the module calculations within the observed variables. These values were calculated from sowing to first flower for each RIL and all sites in the MET dataset, represented by *DayL*_*m*_, *Sradm, Tmax*_*m*_, *and Tmin*_*m*_. Also, *QTL*_*2,g*_, *QTL*_*3,g*_, *QTL*_*5,g*_, *QTL*_*7,g*,_ and *QTL*_*12,g*_ are QTLs that interact with E to affect time to first flower (Bhakta *et al*. 2017). The second part of this equation shows one QTL-by-QTL interaction (*QTL*_*1,g*_ interacting with *QTL*_*12,g*_); *a*_*12*_ is the coefficient for this interaction. The third part of this equation includes the sum of all QTL effects, where *β*_*q*_ represents the coefficient for the *q*^th^ QTL allele effect for RIL_g_ (*QTL*_*q,g*_). Each QTL has a marker value numerically assigned according to its allelic identity; Jamapa alleles were assigned as -1 and Calima alleles as +1 values (see Supporting Information – Table S1).

The daily rates (*FR*_*s,g*_*(t)*) in Eq. (1) are then accumulated or integrated over time to predict day of first flower appearance using Eq. (2) and a daily time step (*dt = 1*).

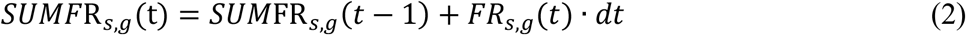

where *SUMFR*_*s,g*_*(t)* integrates the flowering rate at time *t* (in days) starting on the day of planting. *SUMFR*_*s,g*_*(t)* is set to 0.0 at the start of the simulation, and when it reaches 1.0, first flowering is simulated to occur on that day *t* for the *g*^th^ RIL at site *s*.

### 2.4. Dynamic piecewise linear module

The CSM-CROPGRO-Drybean model (CSMG, Table 2), which is part of the Decision Support System for Agrotechnology Transfer (DSSAT; Hoogenboom et al., 2019b), requires daily weather data, soil surface and profile characteristics, crop management scenarios, and cultivar information (GSPs) as input (Jones *et al*. 2003; Hoogenboom *et al*. 2019a). The CSMG crop model uses daily weather variables for maximum and minimum temperature and solar radiation, and these variables have the same units as those used in the DMLM module. However, the day length (h) computed and used in the CSMG model is slightly different from the one used to develop the Bhakta and Vallejos models. The main difference is that the CSMG model accounts for the twilight period at sunrise and sunset, which may affect the photoperiod response of crops. Thus, in this study we used daily day length values computed by the CSMG model as input for the statistical procedures to estimate the numerical coefficients in Eq. (1). This was done to make the daily weather and photoperiod variables identical to those used in the CSMG model (Hoogenboom *et al*. 1994; Boote *et al*. 1998) and to allow incorporation of the new dynamic gene-based module.

A second dynamic module (DMLM-DL) predicts the flowering rate (Eq. (1)) on a daily basis using the daylengths from the CSMG and using a linear response to temperature. However, it is well-known that under high temperature conditions the rate of progression towards flowering does not increase linearly with temperature (White *et al*. 2005). Instead, the response is only approximately linear over a specific range of temperatures, and response plateaus as an optimum temperature is reached. In fact, the effect of temperature on development rate can be more accurately represented by a beta function (Ritchie and Nesmith 1991). Because the temperature varies considerably within a single season, with location, and over time, plants are frequently exposed to temperatures outside their linear response range. To help account for the non-linearity response of common bean under high temperatures, a third module was created that uses a dynamic piecewise linear function (referred to as the DPLM module) to ensure that the daily simulated development rate is bounded to be within the range of temperatures that were observed in the MET study and used to estimate the coefficients in Eq. (1) (Supporting Information – Fig. S2).

*FRMAX*_*g*_ is defined as the rate at which the progress toward flowering proceeds when the environmental conditions, i.e., the daily maximum and minimum temperature, day length, and solar radiation, are at “optimum” values that result in a maximum development rate for the *g*^th^ genotype. The *FRMAX*_*g*_ values were estimated by selecting the maximum rate (1/*DUR*_*s,g*_) for each RIL occurring across *s* environments, using the observed duration between planting and first flower across all *s* sites using Eq. (3):

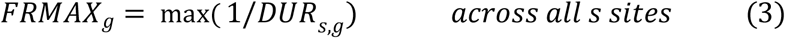

This resulted in 189 data points (one for each RIL plus the two parental lines). We determined whether the values for FRMAXg were affected by the same QTLs that significantly affected the time to first flower by estimating a linear relationship shown in Eq. (4).

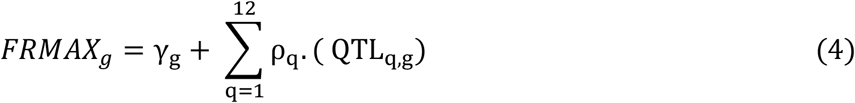

where the variable *γ*_*g*_ is the fixed intercept estimated for the *g*^th^ genotype and *ρ*_*q*_ is the coefficient that quantifies the allelic effect of the *q*^th^ QTL on the maximum rate of progress. A linear regression analysis was used to estimate coefficients of Eq. (4).

The final daily rate of first flowering was determined using the DPLM module that was integrated into the CSMG model in which a maximum rate of development is limited depending on the genotype. If the daily flowering rate at time *t* computed by *FR*_*s,g*_*(t)* exceeds *FRMAX*_*g*_ for any recombinant inbred line in the DPLM module, *FRD*_*s,g*_*(t)* limits the maximum rate of flowering to that set by *FRMAX*_*g*_, Eq. (5):

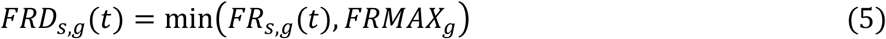

Finally, *FRD*_*s,g*_*(t)* is integrated daily to predict the day when first flower occurs, Eq. (6), where *SUMFRD*_*s,g*_*(t)* integrates the flowering rate at time *t* (in days) in the DPLM module.

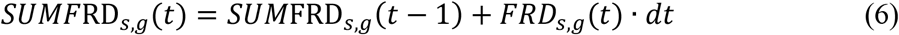

At the start of the simulation, *SUMFRD*_*s,g*_*(t)* is set to 0.0 and the day when it reaches or exceeds 1.0, flowering is predicted to occur for the *g*^th^ genotype.

### 2.5. Incorporation of a gene-based module into CSM

The CSMG model (Hoogenboom *et al*. 1992, 1994) was developed using a modular structure (Jones *et al*. 2001), where overall development and growth are represented by specific modules, including those for vegetative and reproductive development, photosynthesis, respiration, partitioning, vegetative and reproductive growth, and other soil and crop processes (Hoogenboom *et al*. 2019a). For this study the focus was on the phenology module of CROPGRO where the developmental and phenological phase transitions are implemented.

Boote *et al*. (1998) described the physiological development rate in CROPGRO as a function of temperature, photoperiod, and water deficit. If these conditions are optimal, one physiological day is accumulated per calendar day. The phenology module in CROPGRO separates the vegetative and reproductive routines that calculate the stages and individual phase durations.

To incorporate the first flower development stage using *SUMFRD*_*s,g*_*(t)* in the CSMG model, we first developed a new gene-based module (GBM) to create a link between the DPLM module and the crop model (Fig. 4). This module connects daily input data, the DPLM module, and the CSMG phenology module. The inputs for this DPLM module consist of weather data from the crop model and the 12 QTL allelic make up for each RIL (or genotype). The input QTL data for our study were those for the 187 RILs plus the two parent cultivars, which are processed in a new QTL data subroutine inside the GBM module. A new input file was created for the CSMG model, named BNGRO047.GEN that contains QTL data for each of the RILs and their two parent cultivars (Table S1).

The daily weather and QTL data for a particular site and RIL are inputs for the DPLM module, enabling it to simulate the daily flowering rate as affected by G and E conditions. The integrated development progress to first flower, *SUMFRD*_*s,g*_*(t)* and the day when first flowering occurs are passed back to the CSMG phenology routine. The outputs from the GBM module are inputs to the reproductive stage component, where the variables associated with first flowering are calculated. The day when first flowering occurs is set and afterward, progress for subsequent development phases are computed using the original CSMG model.

### 2.6. Sensitivity analysis of simulated variations for G × E

A simulation analysis was performed using the DPLM module to explore all possible combinations of the 12 QTL variations among RILs using the daily weather data across all five sites of the MET study, similar to previous ideotype studies (White *et al*. 2005). This resulted in a total of 4,096 (2^12^) RILs. The coefficients estimated using 187 RILs plus the two parents were used to simulate the number of days to flowering for the 4,096 RILs. The input file BNGRO047.GEN file containing QTL information was revised by adding inputs for each of the 4,096 RILs. Crop management including the planting dates and the daily weather data were assumed to be same as for the original five environments of the MET study. The management input file assumes that only the variations in genetics and environments affect the simulated responses, representing potential production for each line. The DPLM module was then used to conduct the 20,480 unique simulations across sites and synthetic RILs.

The time to first flower responses of the DPLM module were compared with those of the DMLM-DL module to determine how the addition of a maximum rate, *FRMAX*_*g*_ affects the simulated results under different high temperature scenarios in a sensitivity analysis using the 187 RILs and two parents across all five sites of the MET study. The original daily temperature data were used as the base line inputs. Then, both the minimum and maximum temperatures for each day and each site were incremented at a 1 °C increment to create five different temperature scenarios (base, base+1, base+2, base+3, and base+4 C), assuming that crop management, daily solar radiation, and day length were the same as for the original MET study. The simulated number of days to flowering were analyzed and compared for the DPLM and DMLM-DL modules using statistics and a visualization of the distributions of number of days to first. Although we did not have sufficiently high temperatures in the MET study to account for the decrease in development rate as the temperature increases above an optimal threshold, the simple addition of the maximum rate in the piecewise linear module is expected to provide reliable simulations for small increases in temperature.

### 2.7. Yield prediction

An ultimate goal of dynamic crop simulation models is to be able to predict yield. Therefore, we compared the performance of the original CSMG model using GSPs with the DPLM module integrated into CSM-CROPGRO-Drybean model (DPLM-CSM; Table 2). All 18 GSPs were available for 144 genotypes, including 142 RILs and the parent material, except for PA, for which GSPs for only 143 genotypes were available. The procedures for estimating the GSPs were described by Acharya et al. (2017). Daily simulations, starting at planting and continuing until harvest maturity was predicted, were conducted for all five sites for either 2011 or 2012, depending on the MET. Crop management and local weather and soil data based on the original MET study were used as input for the CSMG model (Fig. 1). For the DPLM-CSM hybrid model, the flowering dates were predicted based on the DPLM module using the QTL information as input, rather than the GSPs, while for the other growth and development processes, the GSPs for the individual genotypes were used as input.

### 2.8. Model evaluation

To estimate parameters for the QTL-based modules, we used the *lmer* function of the *lme4* package (Bates *et al*. 2015) of the R programming language (version 3.6.1). To compare the performance of the modules with observed data, we used the estimated parameters for the final QTL-based DPLM module for each site. As a measure of fit of Eq. (1) to the data, we used the root mean squared error (RMSE), defined as

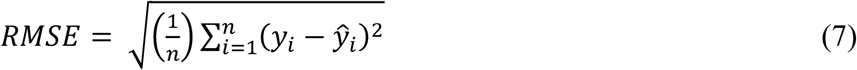

where *y*_*i*_ and *ŷ*_*i*_ are the *i*^th^ observed and simulated number of days to flowering, respectively, and *n* is the number of measurements summed for all values for all RILS and for each site and all sites combined. The adjusted R^2^ was calculated because it indicates module performance adjusted by the number of the terms in the module, defined as

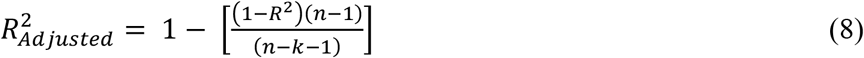

where *R*^*2*^ represents the coefficient of determination, *n* is the number of measurements and *k* is the number of independent variables of the model. A Nash and Sutcliffe (1970) skill score was also used as a measure of model error, referred to as model efficiency (ME) (Wallach *et al*. 2019), and defined as

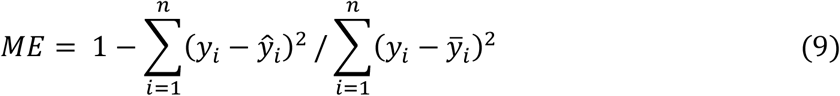

If ME = 1.0, the model fits perfectly, and the observed values are equal to the simulated values (*y*_*i*_ = *ŷ*_*i*_) for each *i* and ME = 1. If ME is less than 0.0, the mean of the observed data is a better predictor of the data than the model. If the variance for the observed minus predicted values is equal to the variance of observations from its mean value, then ME = 0.0, which means that the model is not good because it is no better than using the average of observed values to predict responses. For evaluation of the predictive ability of the modules, we also compared the contributions to prediction error caused by model bias and standard deviation. We used the decomposition of the mean square error (MSE) into bias, standard deviation differences, and residual errors that was developed by Kobayashi and Salam (2000), Eq. (10).

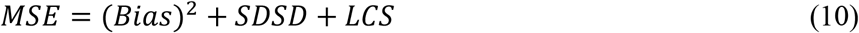

with

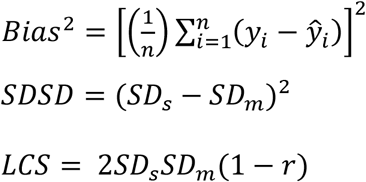

The first term of Eq. (10) is the bias squared, the second term *SDSD* is related to the difference between the simulation standard deviation (SD_s_) and the standard deviation of the measurements (SD_m_). The third term, *LCS*, indicates the remaining MSE error that is not accounted for by bias or standard deviation.

## 3. Results and Discussion

### 3.1. Dynamic mixed linear module coefficients

The *FR*_*s,g*_*(t)* function is shown below with estimated parameter values for the DMLM-DL module. The values of coefficients in Eq. (11) are based on the influence of day length, solar radiation, temperature, and 12 QTL alleles for each RIL, showing G, E, G × E, and G × G interaction effects.

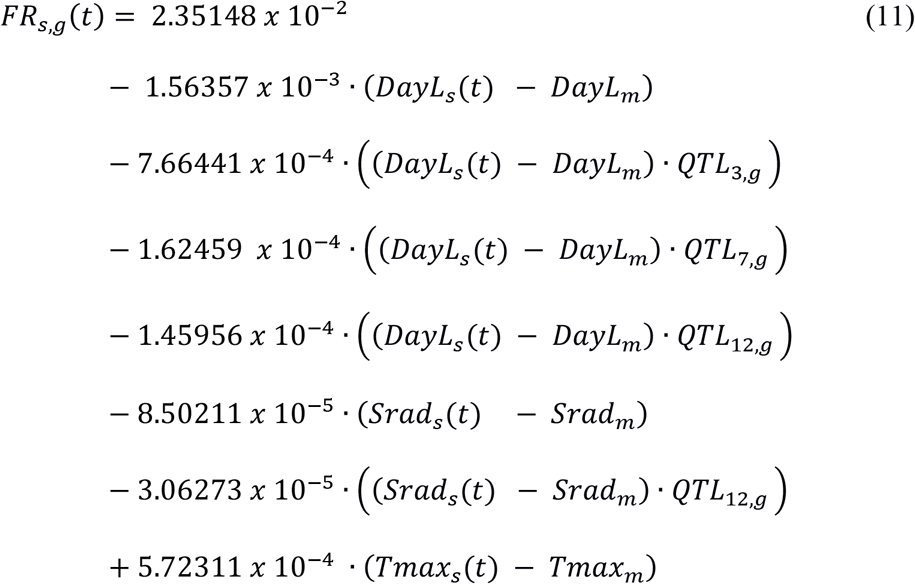

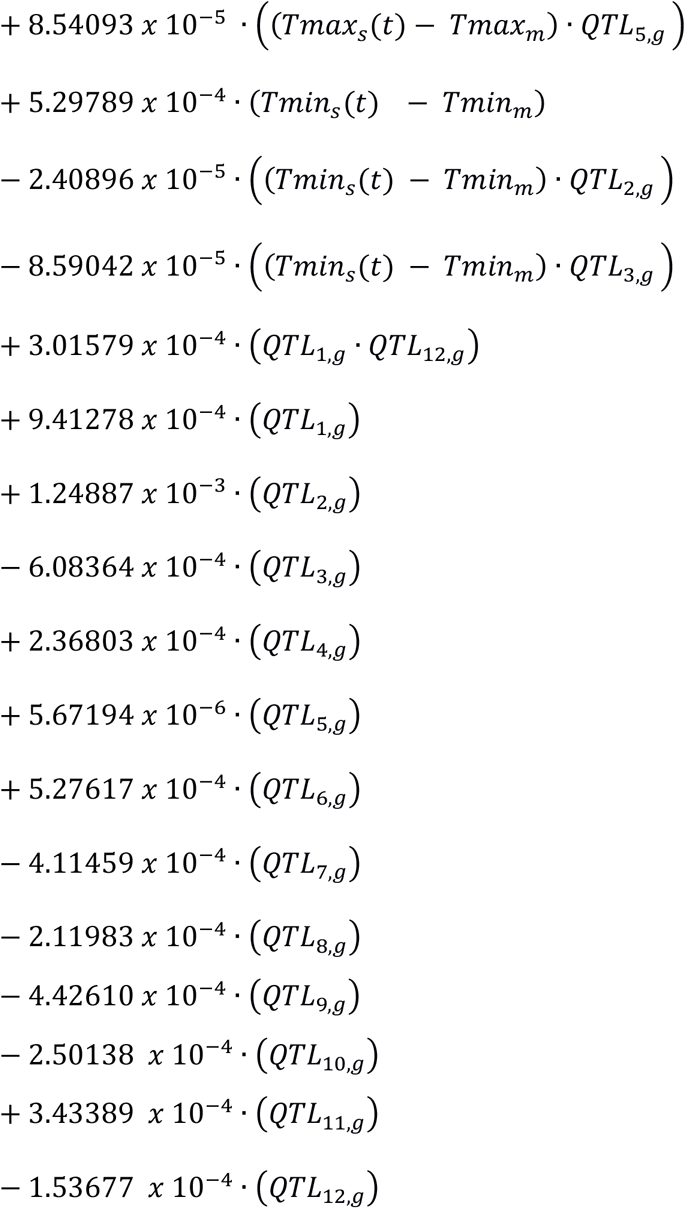

Where the first term (2.35148 × 10^−2^) is the overall average rate of progress, indicating that the average time between sowing and appearance of first flowering is 42.5 days (= 1/(2.35148 × 10^−2^)). The first coefficient (α_1_ = –1.56357 × 10^−3^) is the sensitivity to day length, indicating that a one-hour increase in day length would result in a rate of development that is 1.56357 × 10^−3^ below the average rate of 2.35148 × 10^−2^. This one-hour increase in day length simulates that the time to first flower would occur 45.6 days after planting, an increase of 3.1 days compared to the average days to first flower that was observed across the 5 sites and 187 RILs plus the two parents. This rate of development also varies as a function of QTL alleles, which can increase or decrease the rate resulting in a decrease or increase in the number of days to first flower, respectively. Note that some the QTL coefficients in Eq. (11) have a negative sign while others have a positive sign. This is because each parental genotype has both types of alleles; the allele operator, i.e., Calima = *+1* and Jamapa = -1, will alter the sign of the coefficient accordingly (Table S1). The estimated parameters terms with the 2.5% and 97.5% confidence intervals, p-value, and the variance components are shown in the supporting information (Table S2). The fixed effects variance was 1.80182 × 10^−5^, the random effects variance was 6.34775 × 10^−7^, and the residual variance was 1.3056 × 10^−6^.

Next, we compared the agreement between simulated and observed results for all sites, RILs, and parents using the DMLM and DMLM-DL modules (Fig. 2A, 2B and Table 3). Comparisons of RMSE between simulated and observed values showed that the errors were only slightly different between the two modules (Table 3, DMLM and DMLM-DL modules). When all sites and RILS were included in the comparisons, the RMSE values were 2.73 days and 2.72 days for the DMLM and DMLM-DL, respectively. Similarly, when comparing agreements for each site, the RMSE values using the two module versions were within 0.02 days for ND and 0.04 days for FL. Notably, however, Table 3 shows relatively large differences in RMSE depending on site, with ND having the largest RMSE or 4.58 days in comparison with the lowest RMSE of 1.61 days for PA. We attributed these differences to the fact that the MET did not include a site with long days and low temperatures to contrast the long days and high temperatures of ND, which did not adequately capture the temperature-day length interactions previously documented by Wallace and Enriquez (1980) and Wallace *et al*. (1991).

**Figure 2.**
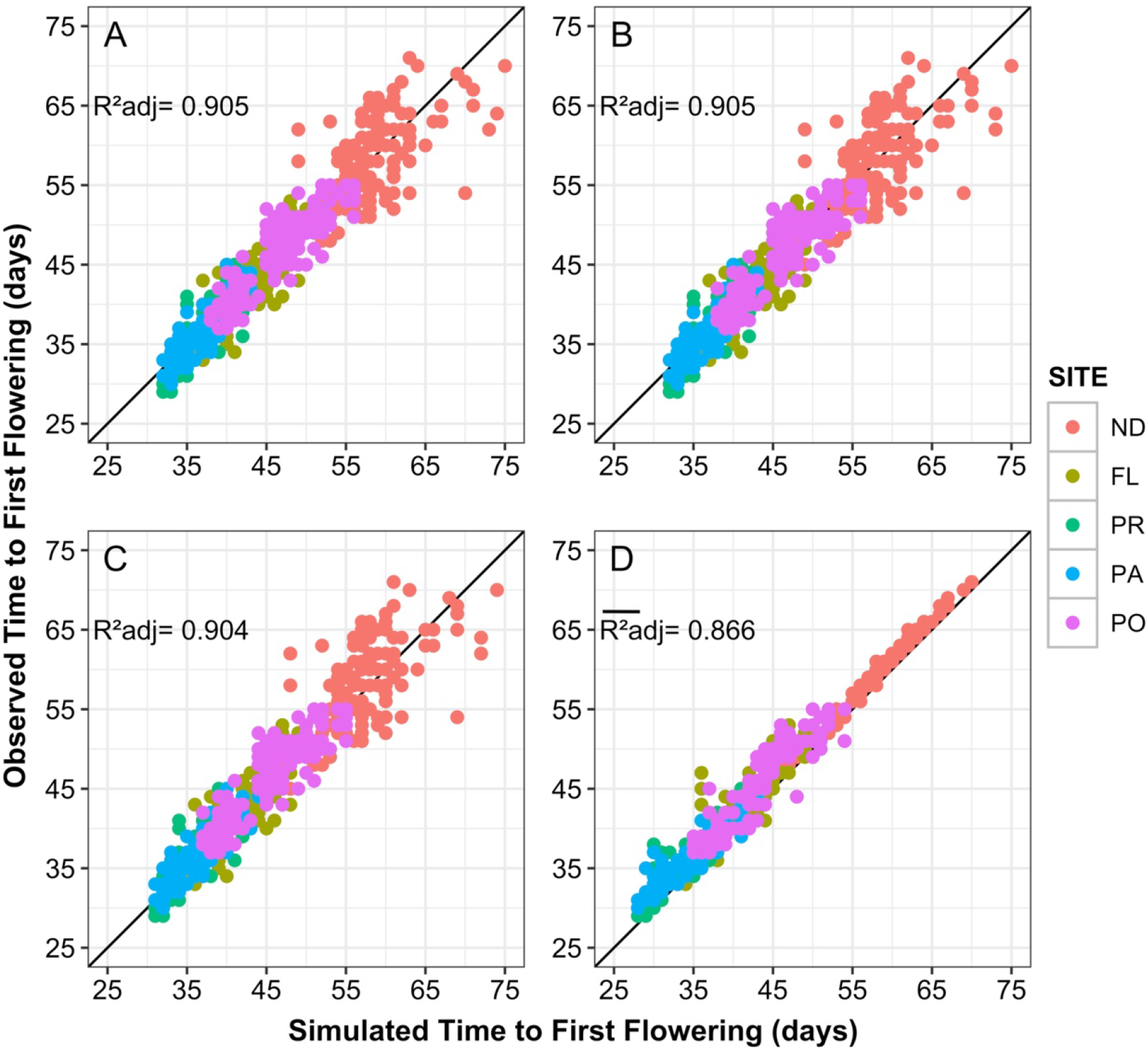
Observed versus simulated time to first flower across all five sites for the dynamic mixed linear module (DMLM) (**A**); the dynamic mixed linear model using the day length computed by the crop module (DMLM-DL) (**B)**; the dynamic piecewise linear module incorporated into CSM-CROPGRO-Drybean (DPLM) (**C**), and the original CSM-CROPGRO-Drybean model using genetic specific coefficients (CSMG) (**D**). For A, B, and C, the modules simulated for each RIL and for all sites, while for D the simulations were conducted based on the genetic specific coefficients based on Acharya *et al*. (2017). Each point represents an observed & simulated RIL; the solid 1:1 diagonal line represents equal values for time to first flower. R^2^adj for graph **D** is the average of the values across all five sites for each RIL. The five experimental sites are: Prosper, North Dakota (ND); Citra, Florida (FL); Puerto Rico (PR); Palmira, Colombia (PA) and Popayan, Colombia (PO).

**Table 3.** Measures of agreement between simulated and observed number of days from planting to first flower for all models for each individual site and for all sites combined.

| Site<sup>1</sup> | Measures of Agreement<sup>2</sup> |  |  |  |  |  |  |  |  |  |
| --- | --- | --- | --- | --- | --- | --- | --- | --- | --- | --- |
|  | Genotypes (#) | Observed Mean | Simulated Mean | Bias | RMSE | ME | MSE | (Bias)<sup>2</sup> | SDSD | LCS |
| **DMLM<sup>3</sup>** |  |  |  |  |  |  |  |  |  |  |
| **ND** | 149 | 57.74 | 58.36 | -0.62 | 4.58 | 0.30 | 21.08 | 0.38 | 0.04746 | 20.65 |
| **FL** | 170 | 42.46 | 42.96 | -0.50 | 2.51 | 0.72 | 6.32 | 0.25 | 1.02153 | 5.05 |
| **PR** | 163 | 36.42 | 36.88 | -0.45 | 1.97 | 0.70 | 3.89 | 0.21 | 0.62889 | 3.05 |
| **PA** | 173 | 36.65 | 37.15 | -0.50 | 1.62 | 0.72 | 2.63 | 0.25 | 0.02306 | 2.36 |
| **PO** | 173 | 45.96 | 46.15 | -0.19 | 2.26 | 0.81 | 5.16 | 0.04 | 0.09782 | 5.02 |
| **All Sites** | 828 | 43.54 | 43.99 | -0.45 | 2.73 | 0.90 | 7.45 | 0.20 | 0.04168 | 7.20 |
| **DMLM-DL** |  |  |  |  |  |  |  |  |  |  |
| **ND** | 149 | 57.74 | 58.32 | -0.57 | 4.56 | 0.30 | 20.90 | 0.33 | 0.09217 | 20.48 |
| **FL** | 170 | 42.46 | 43.03 | -0.56 | 2.55 | 0.71 | 6.53 | 0.32 | 1.11594 | 5.10 |
| **PR** | 163 | 36.42 | 36.88 | -0.45 | 1.97 | 0.70 | 3.89 | 0.21 | 0.58810 | 3.09 |
| **PA** | 173 | 36.65 | 37.18 | -0.53 | 1.61 | 0.73 | 2.61 | 0.28 | 0.01977 | 2.31 |
| **PO** | 173 | 45.96 | 46.08 | -0.12 | 2.23 | 0.81 | 5.02 | 0.01 | 0.10511 | 4.90 |
| **All Sites** | 828 | 43.54 | 43.98 | -0.44 | 2.72 | 0.90 | 7.42 | 0.20 | 0.05778 | 7.17 |
| **DPLM** |  |  |  |  |  |  |  |  |  |  |
| **ND** | 149 | 57.74 | 57.37 | 0.38 | 4.55 | 0.30 | 20.82 | 0.14 | 0.11116 | 20.57 |
| **FL** | 170 | 42.46 | 42.14 | 0.32 | 2.49 | 0.72 | 6.22 | 0.10 | 0.96639 | 5.15 |
| **PR** | 163 | 36.42 | 36.01 | 0.41 | 1.93 | 0.71 | 3.75 | 0.17 | 0.44665 | 3.13 |
| **PA** | 173 | 36.65 | 36.36 | 0.29 | 1.56 | 0.74 | 2.44 | 0.08 | 0.00026 | 2.36 |
| **PO** | 173 | 45.96 | 45.07 | 0.89 | 2.41 | 0.78 | 5.84 | 0.79 | 0.10968 | 4.94 |
| **All Sites** | 828 | 43.54 | 43.08 | 0.46 | 2.73 | 0.90 | 7.46 | 0.21 | 0.06855 | 7.17 |
| **CSMG** |  |  |  |  |  |  |  |  |  |  |
| **ND** | 144 | 57.59 | 56.35 | 1.24 | 1.38 | 0.94 | 1.90 | 1.545 | 0.00366 | 0.35 |
| **FL** | 144 | 41.85 | 40.47 | 1.38 | 2.20 | 0.76 | 4.85 | 1.891 | 0.41912 | 2.54 |
| **PR** | 144 | 36.31 | 34.32 | 1.99 | 2.30 | 0.58 | 5.31 | 3.972 | 0.01996 | 1.32 |
| **PA** | 143 | 36.29 | 34.87 | 1.43 | 1.95 | 0.56 | 3.83 | 2.035 | 0.38940 | 1.41 |
| **PO** | 144 | 45.54 | 43.27 | 2.27 | 3.00 | 0.66 | 9.03 | 5.157 | 0.47913 | 3.40 |
| **All Sites** | 719 | 43.53 | 41.87 | 1.66 | 2.23 | 0.94 | 4.98 | 2.762 | 0.00011 | 2.21 |

<sup>1</sup>Prosper, North Dakota (ND); Citra, Florida (FL); Isabella, Puerto Rico (PR); Palmira, Colombia (PA), and Popayan, Colombia (PO).
<sup>2</sup> ME = Nash and Sutcliffe (1970) model efficiency; MSE = mean squared error; Bias<sup>2</sup>, SDSD, and LCS present the decomposition of MSE.
<sup>3</sup> DMLM = Dynamic Mixed Linear Module; DMLM-DL = Dynamic Mixed Linear Module with day length from the CSM model; DPLM = Dynamic Piecewise Linear Module as integrated in CSM-CROPGRO-Drybean for flowering prediction; CSMG = Original CSM-CROPGRO-Drybean model using GSPs

The comparison of ME between the two module versions (DMLM and DMLM-DL) showed that both have the same high model efficiency value of 0.90. Although ME values for the ND site were lower (0.30 for both modules), these positive numbers indicate that the modules are more effective than using the mean value of the observations. Table 3 also shows that the bias in simulating time to flowering was low across all sites (less than 0.5 days), and MSE values were low except for ND. The remaining error after accounting for bias and standard deviation differences were much larger for both module versions at ND than for any of the other sites. Overall, the module implementation using the CSMG-computed day lengths (DMLM-DL) showed that the agreement indicators were only slightly different from the DMLM module using the day lengths from Bhakta *et al*. (2017).

### 3.2. Dynamic piecewise linear module

The highest maximum rate for any genotype in the MET dataset was 0.0345 and the lowest observed maximum rate for any genotype was 0.0222. This means that the duration from planting to first flower varied from 29 to 45 days among genotypes under optimal environmental conditions. These maximum rates of development occurred at the tropical PA and PR locations where temperatures were warm and day lengths were relatively short. Although environmental conditions may not have been optimal at these locations, most of the maximum rates across locations for any RIL occurred in PA; only a few occurred in PR where the maximum rates for some RILs were only slightly higher than in PA. These results could likely be improved by using other datasets, ideally under more controlled environmental conditions.

The main purpose of Eq. (12) is to prevent predictions of excessively high values for the development rate that could lead to unrealistically low predictions for the number of days to first flower appearance under environmental conditions with a high temperature, a short day length, and a high solar radiation values that are likely to occur in many environments. This equation was incorporated in the DPLM module and integrated into the full DPLM-CSM hybrid model.

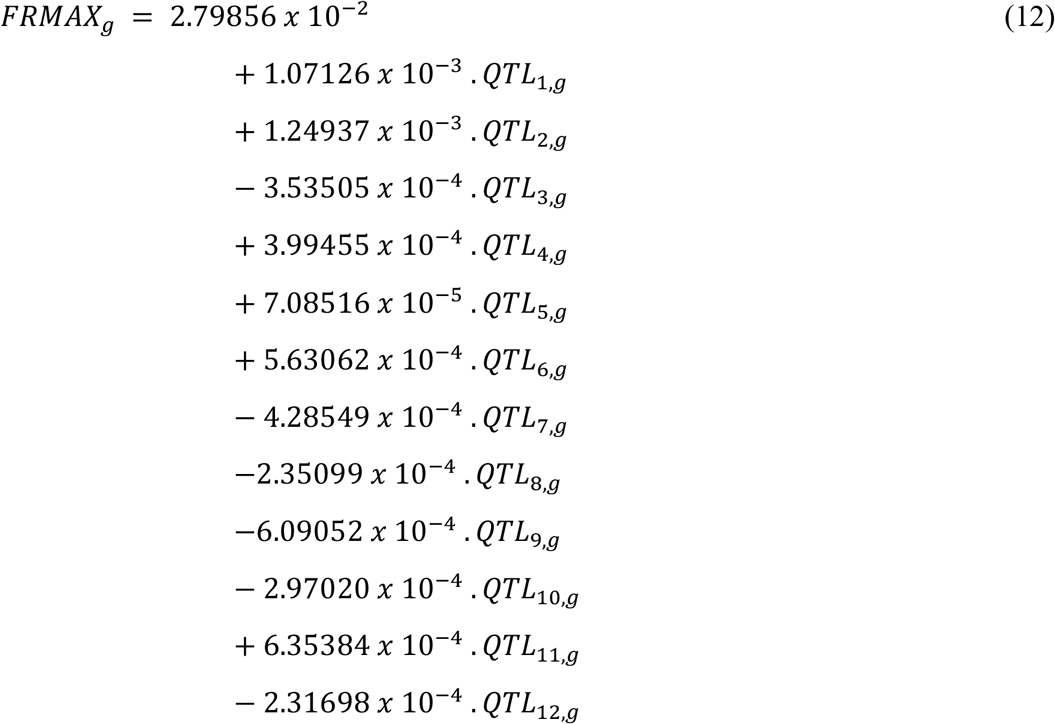

The fitting of *FRMAX*_*g*_ using Eq. (12) resulted in predicted caps on the rate of progress for the 187 RILs plus the two parents of our dataset with a RMSE of 1.66 days, ME of 0.77 days and MSE of 2.78 (Fig. 3). These values indicate that the maximum developmental rates were affected by the genetic factors (12 QTLs).

**Figure 3.**
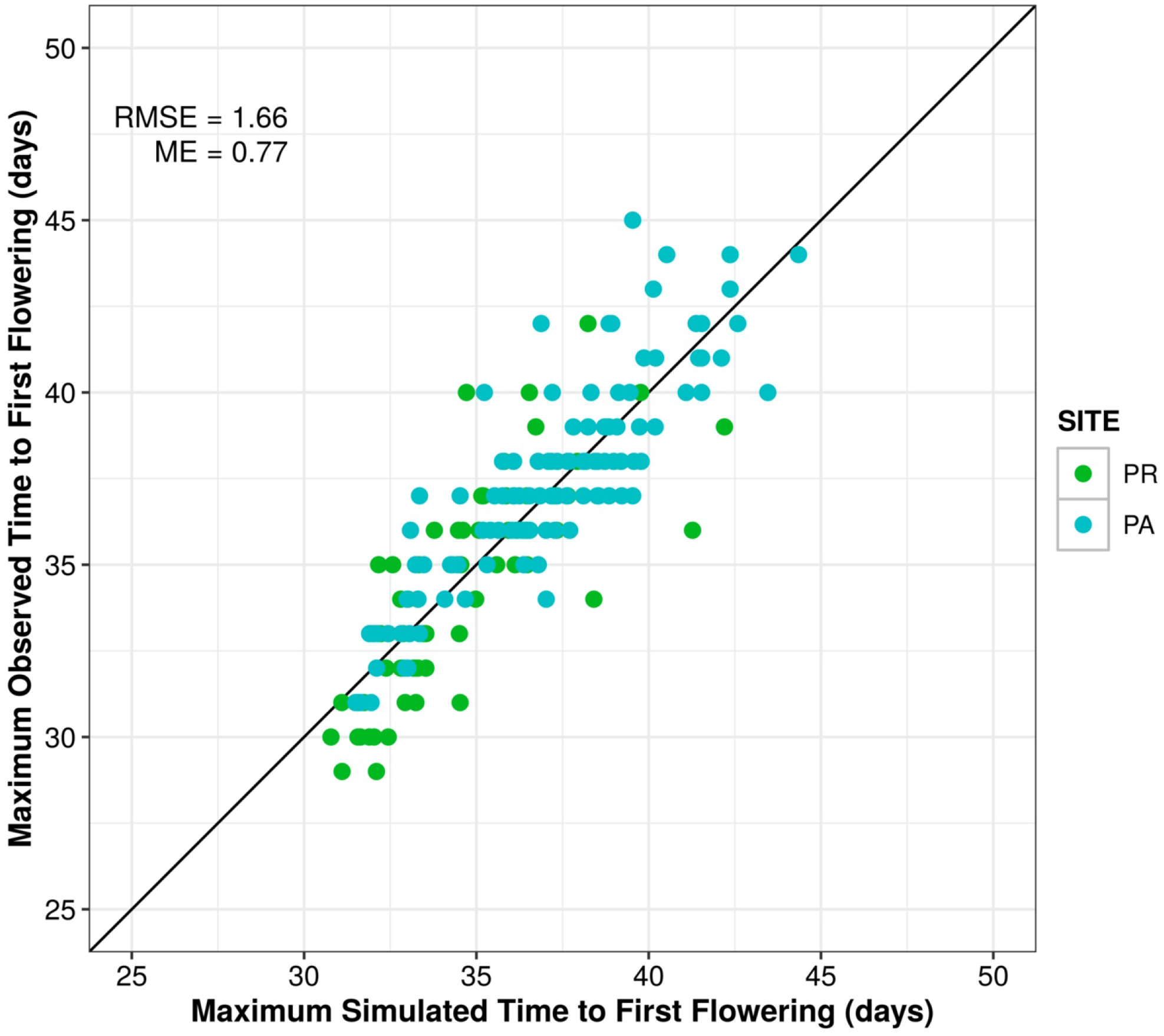
Maximum observed versus simulated time to first flowering for each RIL across all five sites based on a linear model dependent on the 12 QTLs alleles for the 187 RIL plus the two parental lines. RMSE = Root Mean Square Error; ME = Model Efficiency (Nash and Sutcliffe, 1970). The solid 1:1 diagonal line represents equal values of maximum simulated/observed time to first flowering.

**Figure 4.**
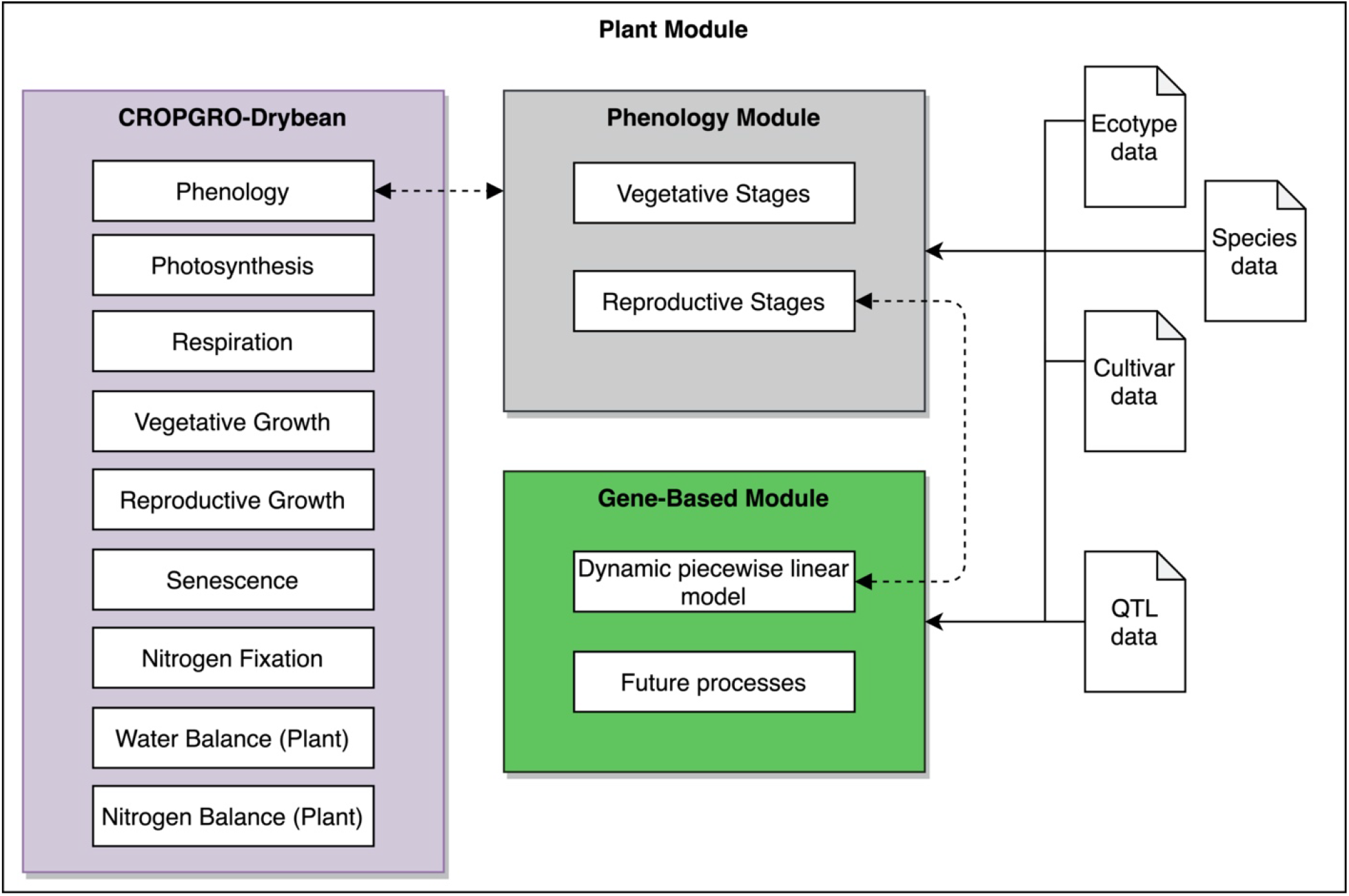
Overview of the DPLM-CSM model developed to integrate the CSM-CROPGRO-Drybean model (CSMG) with the Dynamic Piecewise Linear Module (DPLM) using a new gene-based module (GBM). The DPLM simulates the first time of first flowering module developed from the dynamic mixed linear model first developed by Vallejos *et al*. (2020). The integrated model uses QTL data, which contains the 12 QTL allele information to simulate the daily rate of development towards first flowering, in addition to the other input data used by the original CSMG.

### 3.3. Structural changes of the CSM-CROPGRO-Drybean model

The new gene-based module (GBM) operates on a daily time step in the DPLM-CSM phenology module to simulate the rate of development towards first flowering for a particular RIL or cultivar and for a specific site, as shown in Fig. 4. This GBM module incorporates the DPLM module and processes the QTL data obtained from the revised BNGRO047.GEN file, while weather data are passed to the DPLM module from the CSMG routines. When the value of *SUMFRD*_*s,g*_*(t)* reaches 1.0 (Eq. (6)), the day of first flower is simulated to occur and this date is passed back to the phenology module for its use in updating first flowering in the reproductive development module.

The DPLM module was designed to be flexible, operating in parallel with the original CSMG using GSPs. This allows the CSMG model to work in a hybrid mode using either the original cultivar coefficients or the QTL input data to simulate the development of first flowering. Regardless, all other stages in the DPLM-CSM model are simulated using inputs from the original cultivar coefficient file. This option was added as a new switch in the crop management input file (FileX). When this switch is set to ‘Y’ the DPLM-CSM model uses the DPLM module and the QTL input data to simulate the time of first flower. Otherwise if the switch is set to ‘N’, the DPLM-CSM model uses the original GSPs for all phenological development stages, including the prediction of flowering, and the DPLM module is ignored. These changes do not affect any other phenological processes in the crop growth model. In this way, additional dynamic gene-based modules can be added to the GBM to simulate other vegetative and reproductive processes.

### 3.4. Comparing simulated and observed frequency distributions of time to flower

The simulated and observed frequency distributions of days between sowing and first flower are presented in Fig. 5 for the DPLM module simulations (left panel) and for observed data from the MET study (right panel). The shapes of the simulated distributions appeared to be bimodal for all locations except for ND where it showed a distribution close to normal. The distributions for the observed data did not exhibit bimodal characteristics, except for the PO site, which had cooler temperatures than the other sites. QTL2, which is associated with the growth habit gene *Fin*, shows interaction with Tmin and is likely responsible for this bimodality (Bhakta *et al*. 2017). Also, on average, the indeterminate growth habit RILs generally flowered later than the determinate growth habit RILs. These graphs showed that Calima flowered earlier than Jamapa except for the ND site. The time-to-flowering pattern of the two parents was captured by the module. Bhakta et al. (2017) detected this transgressive behavior of some RILs, those flowering earlier or later than the parents, a phenomenon explained by the presence of genes that accelerate development and others that retard development in both parents.

**Figure 5.**
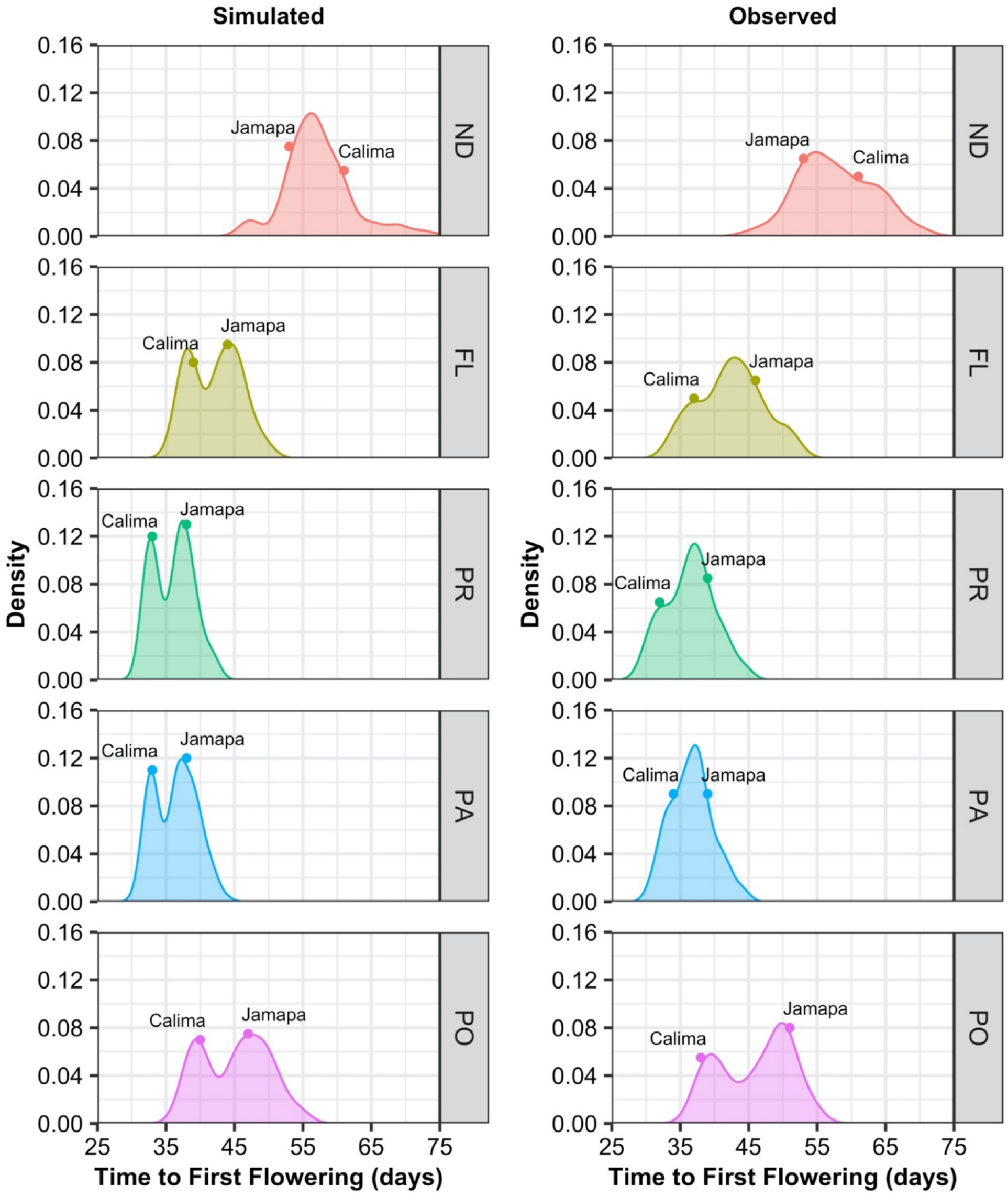
Density plots of time to first flower in days across five sites. Distribution of simulated time to first flower using the dynamic piecewise linear module (DPLM) (left panel) and the distribution of observed time to first flower (right panel). The parental lines Jamapa and Calima are highlighted at the top of each distribution. The five experimental sites are: Prosper, North Dakota (ND); Citra, Florida (FL); Puerto Rico (PR); Palmira, Colombia (PA) and Popayan, Colombia (PO).

Table 3 shows a comparison using various measures of agreement between the simulated and observed data for DMLM, DMLM-DL, DPLM, and CSMG. The simulation results of the DPLM module displayed a strong agreement between the simulated and observed time to first flower (Fig. 2C and Table 3). Simulated results showed an average bias of 0.55, a RMSE of 2.73 days, a ME of 0.90, a MSE of 7.46 and an adjusted R^2^ of 0.905. The differences between the simulated and observed values were larger for ND than for the other sites. The average bias for ND was 0.38, the RMSE was 4.55 days, and the ME was 0.30, whereas the corresponding values for the other four sites showed a much closer agreement between the simulated and observed days to first flower. Comparisons of these agreement indicators with those for the DMLM and DMLM-DL modules showed nearly identical bias, RMSE, and ME values, demonstrating that the implementation of the DPLM module provided simulated results that were nearly identical to the other two module versions listed in Table 3.

The original CSMG mostly produced simulated days to first flower that were in closer agreement with observed results across all sites than the other modules (Table 3 and Fig. 2D). However, these results are misleading in that the GSPs that produced these results were estimated for each individual RIL, which means that only 5 data points were used to estimate 3 GSPs for each RIL, and thus the agreements were forced in the GSP estimation process. The adjusted R^2^ was calculated using five parameters for each RIL; the 113 RILs that had observations for all five sites resulted in estimating 339 GSP parameters. The adjusted R^2^ averaged for the RILs was 0.866, ranging from 0.321 to 0.997 with a standard deviation of 0.129. However, note that the adjusted R^2^ values were lower than all of those for the gene-based modules for each site except at ND. As Acharya et al. (2017) point out, the estimated GSPs were highly uncertain and that different combinations of the GSPs could provide the same fit to observed data (showing equifinality in the estimation process) such that the GSP estimates are not reliable even though they can nearly reproduce the data. Estimating three parameters with only five data points, then repeating this process for each of the RILs, results in estimates that reliably reproduce the data used to estimate them but should not be interpreted as values that can be used for other environments or genotypes. Estimation of these coefficients also requires considerable effort and resources, which has to be repeated every time a new cultivar is released. Instead statistical gene-based modules can estimate independently phenotypic traits using as input G, E, and G × E interactions data. By contrast, estimating the 25 coefficients in the dynamic linear module (Eq. (11)) used all data across the five sites and 189 (RILs plus parents), thus 945 observations were used to estimate 25 coefficients. Therefore, using the dynamic mixed linear module estimation process has potential for a more robust use of the module across environments and genotypes, especially for a new genotype that has QTL information but does not have field phenotype data.

The frequency distributions associated with genetic variations in the RIL population for simulated time to first flower at each site and for the five temperature scenarios are shown in Fig. 6. The comparisons of the means and standard deviations of the populations for each 4-temperature/site combination are summarized in Table 4. These results demonstrate the effects of including the maximum rate of development (FRMAX_g_) for each genotype in the DPLM module for comparison with the module without this upper limit. The largest differences in simulated days to first flower occurred when the temperature was increased by 4 °C (Table 4) at sites with higher temperatures. For example, for the PR site, increasing the daily Tmin and Tmax values by 4 °C only decreased the mean days to first flower by 0.6 days for the DPLM module, whereas the increase in temperature by 4 °C unrealistically decreased the mean days to first flower by 5.1 days for the DMLM-DL module. In contrast, results for the cooler sites (PO and ND) were similar for both module versions. The frequency distributions (Fig. 6) visually demonstrate the effect of the FRMAX_g_ on days to first flower. The distributions for PO and ND shifted to the left for each temperature increase of 1 °C, indicating a more rapid rate of development for each site, whereas the distributions of responses of the same populations at the warm sites (PR and PA) changed very little even for the 4 °C temperature increase.

**Figure 6.**
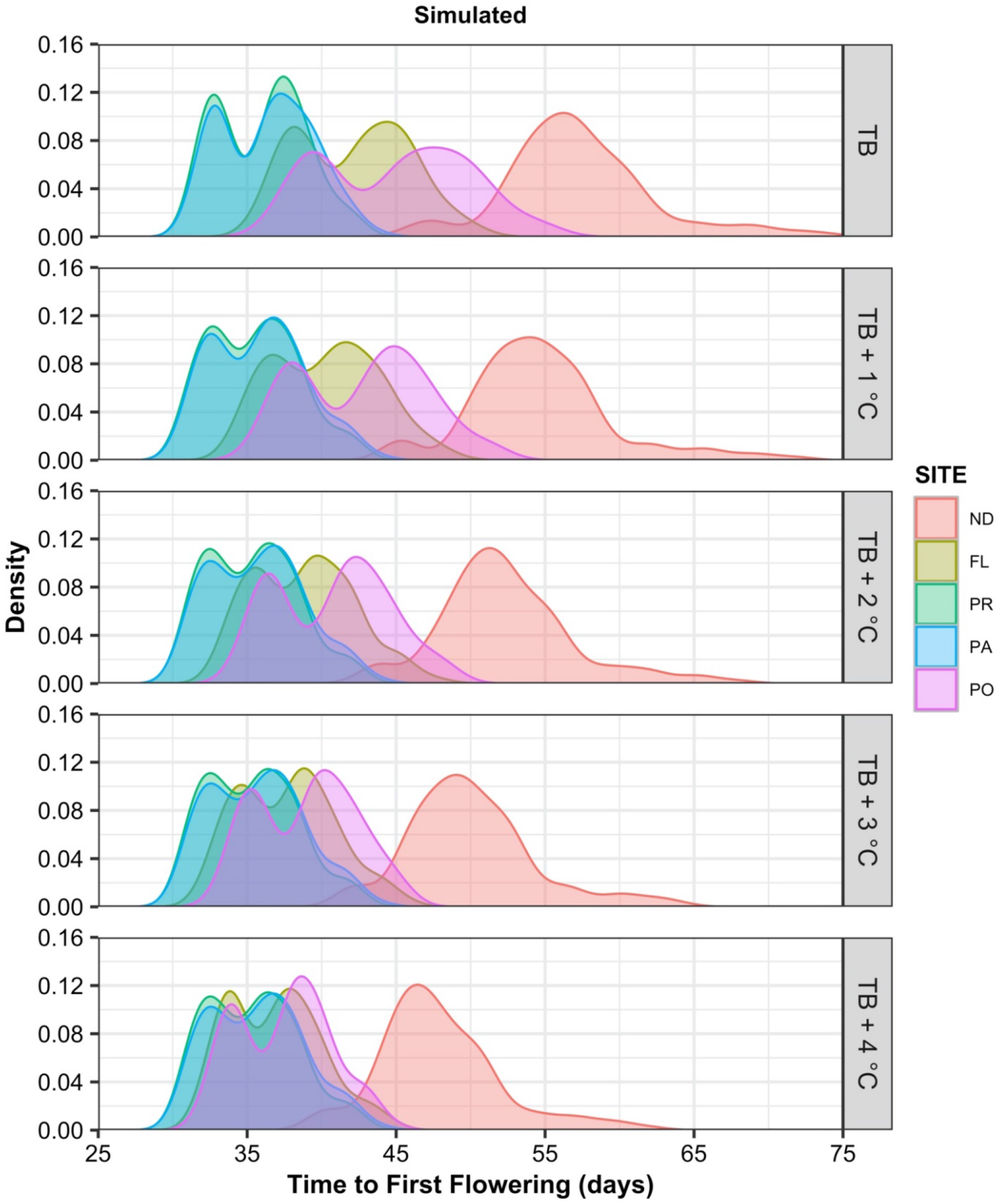
Density plots of distributions for simulated time to first flower (in days) using the dynamic piecewise linear module (DPLM). Simulated days between planting to first flower shows the responses to increasing the base maximum and minimum temperature from 1 °C through 4 °C. The five experimental sites are: Prosper, North Dakota (ND); Citra, Florida (FL); Puerto Rico (PR); Palmira, Colombia (PA) and Popayan, Colombia (PO).

**Table 4.** Temperature *s*ensitivity analysis for the simulated number of days to first flower for the dynamic piecewise linear module (DPLM) and the dynamic mixed linear module with CSM-CROPGRO-Drybean day length (DMLM-DL) using the original weather data from the five sites

| Site<sup>1</sup> | DPLM |  |  |  | DMLM-DL |  |  |  |
| --- | --- | --- | --- | --- | --- | --- | --- | --- |
|  | Simulated Mean | Min<sup>2</sup> | Max<sup>2</sup> | Standard Deviation | Simulated Mean | Min | Max | Standard Deviation |
| **Base Temperature** |  |  |  |  |  |  |  |  |
| **ND<sup>1</sup>** | 57.37 | 46 | 76 | 5.14 | 58.32 | 47 | 77 | 5.17 |
| **FL** | 42.14 | 35 | 51 | 3.74 | 43.03 | 36 | 51 | 3.67 |
| **PR** | 36.01 | 31 | 43 | 2.94 | 36.88 | 32 | 43 | 2.85 |
| **PA** | 36.36 | 31 | 44 | 3.07 | 37.18 | 32 | 43 | 2.94 |
| **PO** | 45.07 | 37 | 55 | 4.87 | 46.08 | 38 | 56 | 4.88 |
| **All Sites** | 43.08 | 31 | 76 | 8.56 | 43.98 | 32 | 77 | 8.58 |
| **Base Temperature + 1 °C** |  |  |  |  |  |  |  |  |
| **ND** | 54.74 | 45 | 72 | 4.81 | 55.66 | 46 | 73 | 4.83 |
| **FL** | 40.29 | 34 | 49 | 3.53 | 41.04 | 35 | 49 | 3.38 |
| **PR** | 35.48 | 30 | 43 | 2.97 | 35.40 | 31 | 41 | 2.55 |
| **PA** | 35.77 | 30 | 44 | 3.10 | 35.69 | 31 | 41 | 2.65 |
| **PO** | 42.87 | 36 | 52 | 4.20 | 43.86 | 37 | 53 | 4.21 |
| **All Sites** | 41.54 | 30 | 72 | 7.77 | 42.03 | 31 | 73 | 8.02 |
| **Base Temperature + 2 °C** |  |  |  |  |  |  |  |  |
| **ND** | 52.37 | 43 | 68 | 4.49 | 53.13 | 44 | 69 | 4.58 |
| **FL** | 38.81 | 33 | 48 | 3.36 | 39.29 | 34 | 47 | 3.07 |
| **PR** | 35.39 | 30 | 43 | 2.99 | 34.09 | 30 | 39 | 2.33 |
| **PA** | 35.69 | 30 | 44 | 3.13 | 34.29 | 30 | 40 | 2.41 |
| **PO** | 40.81 | 34 | 49 | 3.78 | 41.80 | 35 | 50 | 3.78 |
| **All Sites** | 40.34 | 30 | 68 | 6.97 | 40.24 | 30 | 69 | 7.51 |
| **Base Temperature + 3 °C** |  |  |  |  |  |  |  |  |
| **ND** | 50.03 | 42 | 64 | 4.26 | 50.70 | 42 | 65 | 4.39 |
| **FL** | 37.67 | 32 | 46 | 3.21 | 37.68 | 33 | 45 | 2.71 |
| **PR** | 35.36 | 30 | 43 | 3.00 | 32.85 | 29 | 38 | 2.14 |
| **PA** | 35.68 | 30 | 44 | 3.13 | 33.07 | 29 | 38 | 2.20 |
| **PO** | 39.03 | 33 | 46 | 3.31 | 40.01 | 34 | 47 | 3.30 |
| **All Sites** | 39.31 | 30 | 64 | 6.21 | 38.60 | 29 | 65 | 7.00 |
| **Base Temperature + 4 °C** |  |  |  |  |  |  |  |  |
| **ND** | 47.91 | 40 | 62 | 4.08 | 48.48 | 41 | 63 | 4.14 |
| **FL** | 36.88 | 32 | 45 | 3.06 | 36.21 | 32 | 43 | 2.46 |
| **PR** | 35.36 | 30 | 43 | 3.00 | 31.77 | 28 | 36 | 1.96 |
| **PA** | 35.68 | 30 | 44 | 3.13 | 31.95 | 28 | 37 | 2.01 |
| **PO** | 37.50 | 32 | 45 | 3.12 | 38.34 | 33 | 45 | 3.01 |
| **All Sites** | 38.44 | 30 | 62 | 5.57 | 37.10 | 28 | 63 | 6.54 |

<sup>1</sup>Prosper, North Dakota (ND); Citra, Florida (FL); Isabella, Puerto Rico (PR); Palmira, Colombia (PA), and Popayan, Colombia (PO).
<sup>2</sup> Minimum/Maximum simulated number of days from sowing to first flower.

We are not suggesting that use of the upper limit on development rate is robust for broad use, but instead that the MET should include more sites that have a wider range of temperatures and day lengths to enable nonlinear responses to be estimated. For example, improvements could be attained using a beta function for temperature response, in addition to controlled environment experiments with a wider range of genetic material to develop nonlinear functions to represent the full range of environmental responses in this crop species.

### 3.5. Simulating response distributions for all potential genotype combinations

The performance of DPLM module across the five experimental sites was simulated for RILs with all possible allelic combination (4096) of the twelve QTLs used by the dynamic time-to flower module. Frequency distributions for the number of days from planting to first flower were produced for each location (Fig. 7). The dots in the figure highlight the number of days required for the Jamapa and Calima parents to reach the stage of first flowering. The simulated first flowering dates at the ND site were later (mean of 59.1 days) and had a larger standard deviation (9.10 days) compared to the other sites. The spread of simulated days to first flower ranged between 41 and 94 days at ND due to its longer day lengths and some days with cooler temperatures than other sites. The smallest average number of days to first flower was for sites with high temperature conditions and short day lengths (PR and PA), where simulated means were about 36 days for both locations, and standard deviations of 2.6 and 2.6 days, respectively, with response ranges varying from 34 to 39 days for each location. The FL and PO sites with their warm conditions showed simulated means of 42 days and 45 days, respectively, and standard deviations of 3.5 days and 3.9 days, respectively. The shapes of the distributions were bell-shaped across all sites except for ND which showed the flattest shape due to the G, E, G × E, and G × G interactions in the mixed piecewise dynamic module. The altered behavior of the parental lines was also found in these simulations, where Jamapa flowered earlier than Calima for the FL, PA, PO, and PR sites and Calima flowered earlier than Jamapa for the ND site.

**Figure 7.**
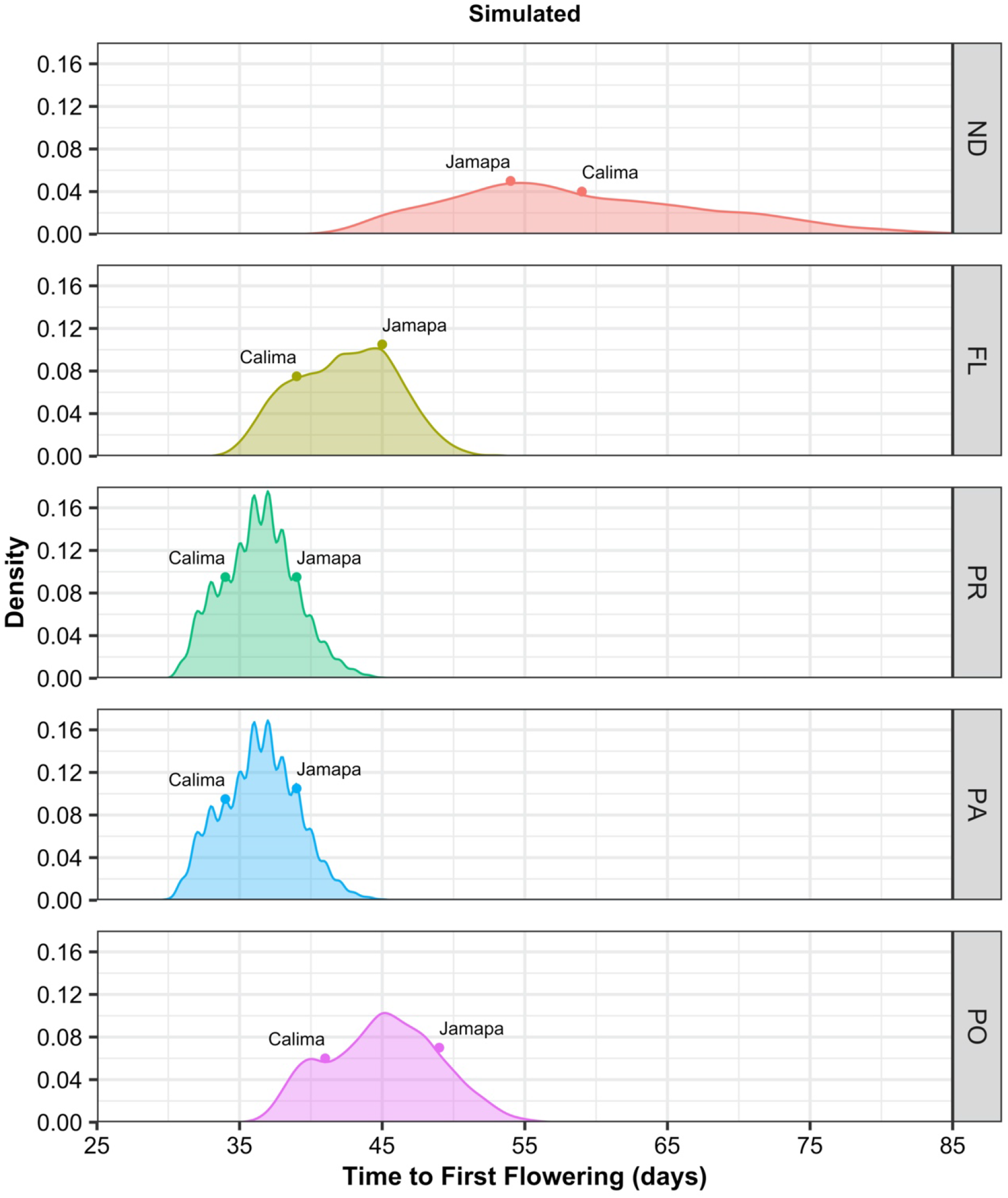
Density plots for simulated time to first flower (days) across the five sites showing all possible genetic combinations. The distribution of simulated days to flower by site includes all recombinant inbred line combinations (2^12^ = 4096) as simulated by the dynamic piecewise linear module (DPLM), while dots at the top of each distribution represent the simulated parental lines Jamapa and Calima. The five experimental sites are: Prosper, North Dakota (ND); Citra, Florida (FL); Puerto Rico (PR); Palmira, Colombia (PA) and Popayan, Colombia (PO).

### 3.6. Yield prediction

For each of the five sites, we simulated yield using the original CSMG model and the original GSPs that were calibrated by Acharya et al. (2017) for each individual RIL. For the MET crop management practices and one year of environmental conditions (Fig 1), simulated mean yield by CSMG was lowest for PA (190.0 ± 89 kg/ha) and highest for ND (637.3 ± 242 kg/ha) while for the other three sites mean yield ranged from 304.7 ± 135 kg/ha for FL, 505.0 ± 270.0 kg/ha for PO, and 540.0 ± 244.8 kg/ha for PR (Fig. 8). For the DPLM-CSM hybrid model, simulated yield ranking among the five sites was similar. The highest mean yield was obtained for ND (720.0 ± 291.0 kg/ha), while the lowest mean yield was obtained for PA (234.4 ± 104.2 kg/ha). Mean yield for FL was 354.8 ± 156.4 kg/ha, for PO was 625.0 ± 311.2 kg/ha, and for PR was 673.3 ± 264.7 kg/ha (Fig. 8). The differences in yield were due to the slight differences in simulated flowering dates between the original CSMG model and the DPLM-CSM hybrid model (Fig.2 and 5), while all other growth and development processes were simulated exactly the same as the CSMG model using the same inputs (Fig. 4).

**Figure 8.**
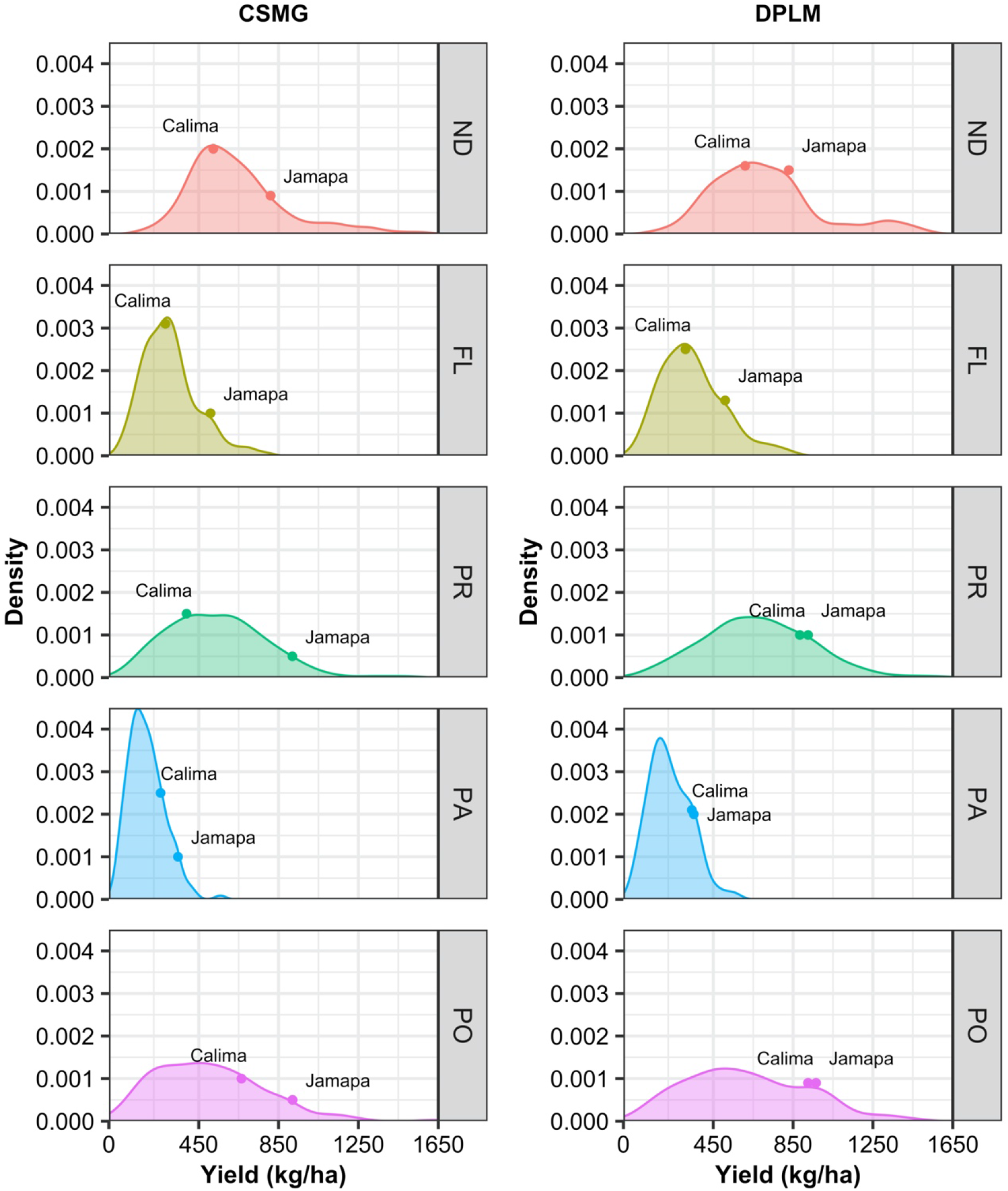
Density plot for simulated yield using the original CSM-CROPGRO-Drybean model and genetic specific coefficients (CSMG) (left panel) and the dynamic piecewise linear module (DPLM) integrated with the CSM-CROPGRO-Drybean model (right panel). The five experimental sites are: Prosper, North Dakota (ND); Citra, Florida (FL); Puerto Rico (PR); Palmira, Colombia (PA) and Popayan, Colombia (PO).

### 3.7. Further advancement in gene-based modeling

This work presents an approach for incorporating gene-based modules into an existing crop growth model for simulating days to first flower (by the DPLM module) and simulating all other processes and final yield using original components of the CSMG. It builds on the approach discussed by Vallejos *et al*. (2020). Only minor modifications were needed to enable their dynamic model to be integrated as a module into the existing CSMG model (Hoogenboom *et al*. 1994; Jones *et al*. 2003). There were only small differences between results from the dynamic piecewise linear module integrated into the CSMG model from our work and model results published by Vallejos *et al*. (2020). We recognize the need for use of independent data to evaluate the predictive capabilities in other environments and are working on that. In addition, there is a need to use a more diverse population to evaluate the ability of the model to predict first flower occurrence across genetic variations that may not be in the population used in the MET dataset.

The model integration approach used here is different from previously published approaches because it incorporates a gene-based dynamic model to replace an existing dynamic component in a comprehensive crop growth simulation model. The approach used for integrating the gene-based first flower module into the CSMG model possibly can be used to incorporate other gene-based modules to systematically transition from a GSP-based model to a gene-based model (Hoogenboom *et al*. 2004). In this work, we only added one type of input data, the genetic information for each RIL and parent. All of the other inputs in the original crop model were not modified, and information on planting date and daily weather data were used by the new gene-based time to flower module, ensuring consistency in inputs across all existing and new components of the model.

This work also shows that integrating the genetic information is a promising approach to predict plant development stages of new genotypes and new environments. Instead of estimating GSPs for a specific trait, it requires less effort when a new cultivar is released in that only QTL information is required, saving time and resources that would be otherwise needed for phenotyping. The long-term expectation associated with most QTL studies is the replacement of each QTL linked marker with the gene responsible for that particular QTL effect. This work further shows that genetic modules for other processes can be based on statistical methods that are routinely used by geneticists, if they are developed to replace equivalent modules in existing dynamic crop models.

However, it is clear that considerably more progress is needed to identify other issues that might occur by combining these two types of models. There is a need to extend gene-based modules to cover the full genetic variability of a crop and to introduce other process modules into existing models. Further work is required to improve the gene-based module and to add other processes that are linked dynamically with the crop model.

## 4. Conclusions

This study showed the potential for integrating a process-oriented gene-based module that only requires genetic input information into an existing comprehensive crop model with its empirical cultivar inputs without changing other modules or inputs. The CSM-CROPGRO-Drybean model with the integrated gene-based module was able to not only predict flowering date using only QTL and weather information, but also final yield using the original GSPs for all processes except rate of progression to first flower. This approach can be extended to other processes for which QTL information is readily available.

## Supporting Information

The following additional information is available in the online version of this article — **Table S1:** Recombinant inbred lines for common bean (*Phaseolus vulgaris* L.). Each Quantitative trait loci (QTL) has a marker value according to its allelic identity, assigned as “+1” for the Calima alleles and “-1” for the Jamapa alleles. This information was used as input for the Dynamic Piecewise Linear Module (DPLM) coupled with the CSM-CROPGRO-Drybean model.

**Table S2:** Estimated values in the dynamic QTL effect module showing the estimated parameters with confidence intervals and p-values for the rate of progress from planting to flowering.

**Figure S1:** Parameter estimation process for predicting first flowering across all sites using the Dynamic Mixed Linear Module (DMLM).

**Figure S2:** Computer code for the Dynamic Piecewise Linear Module (DPLM) coupled with the CSM-CROPGRO-Drybean model.

## Acknowledgments

The first author would like to thank the Department of Agricultural and Biological Engineering at University of Florida and the Graduate Program in Applied Computing at University of Passo Fundo for supporting this research.

## Contributions by the authors

F.A.A.O. software development, evaluation, writing; J.W.J. conceptualization, supervision and methodology, formulation and original code of the dynamic mixed linear model; W.P. software development review, supervision; M.B. environmental and QTL data, development of the mixed linear model; C.E.V. development of the genetic material and mixed linear model, formulation and original code of the dynamic mixed linear model; M.J.C. and K.J.B genetic specific coefficients data for CSM-CROPGRO-Drybean model; C.A.H. R programming supervision; J.M.C.F. statistical analysis review; G.H. conceptualization, supervision, writing, and methodology. All co-authors critically reviewed the manuscript.

