## Supplementary Material for "INCORPORATING A DYNAMIC GENE-BASED PROCESS MODULE INTO A CROP SIMULATION MODEL"

***Table S1*** Recombinant inbred lines for common bean (*Phaseolus vulgaris* L.). Each Quantitative trait loci (QTL) has a marker value according to its allelic identity, assigned as “+1” for Calima alleles and “-1” for Jamapa alleles. This information was used as input for the Gene Based Module coupled with the CSM-CROPGRO-Drybean model.

| RIL | QTL1 | QTL2 | QTL3 | QTL4 | QTL5 | QTL6 | QTL7 | QTL8 | QTL9 | QTL10 | QTL11 | QTL12 |
|---|---|---|---|---|---|---|---|---|---|---|---|---|
| Calima | 1 | 1 | 1 | 1 | 1 | 1 | 1 | 1 | 1 | 1 | 1 | 1 |
| Jamapa | -1 | -1 | -1 | -1 | -1 | -1 | -1 | -1 | -1 | -1 | -1 | -1 |
| RIJC001 | -1 | -1 | -1 | 1 | -1 | -1 | -1 | -1 | 1 | 1 | 1 | 1 |
| RIJC002 | 1 | -1 | 1 | 1 | 1 | -1 | -1 | -1 | -1 | 1 | -1 | -1 |
| RIJC003 | 1 | 1 | 1 | 1 | 1 | 1 | -1 | 1 | -1 | -1 | 1 | 1 |
| RIJC004 | -1 | -1 | -1 | -1 | 1 | 1 | -1 | 1 | 1 | 1 | -1 | -1 |
| RIJC005 | 1 | 1 | 1 | -1 | 1 | -1 | -1 | -1 | -1 | -1 | 1 | 1 |
| RIJC006 | 1 | 1 | 1 | 1 | 1 | 1 | -1 | -1 | 1 | -1 | 1 | 1 |
| RIJC007 | 1 | -1 | 1 | 1 | 1 | 1 | -1 | 1 | 1 | -1 | -1 | -1 |
| RIJC008 | 1 | 1 | 1 | 1 | -1 | -1 | -1 | 1 | -1 | -1 | 1 | 1 |
| RIJC009 | -1 | -1 | -1 | 1 | 1 | -1 | -1 | 1 | -1 | 1 | 1 | 1 |
| RIJC011 | -1 | -1 | -1 | -1 | -1 | -1 | -1 | 1 | 1 | 1 | 1 | 1 |
| RIJC012 | 1 | 1 | 1 | 1 | -1 | -1 | -1 | 1 | -1 | -1 | 1 | 1 |
| RIJC013 | 1 | 1 | 1 | 1 | -1 | -1 | -1 | -1 | 1 | -1 | 1 | 1 |
| RIJC014 | 1 | 1 | 1 | 1 | 1 | 1 | 1 | -1 | 1 | 1 | 1 | 1 |
| RIJC015 | 1 | 1 | 1 | 1 | -1 | 1 | -1 | -1 | 1 | -1 | 1 | 1 |
| RIJC016 | -1 | -1 | -1 | -1 | -1 | -1 | 1 | -1 | 1 | -1 | -1 | -1 |
| RIJC017 | -1 | -1 | 1 | 1 | -1 | -1 | -1 | 1 | -1 | -1 | 1 | 1 |
| RIJC018 | 1 | 1 | 1 | 1 | 1 | 1 | 1 | -1 | 1 | 1 | -1 | 1 |
| RIJC019 | 1 | 1 | 1 | 1 | -1 | -1 | 1 | 1 | -1 | 1 | -1 | -1 |
| RIJC020 | 1 | 1 | -1 | -1 | -1 | -1 | -1 | -1 | 1 | 1 | -1 | -1 |
| RIJC021 | 1 | 1 | 1 | -1 | -1 | 1 | 1 | 1 | -1 | -1 | -1 | -1 |
| RIJC022 | -1 | -1 | 1 | 1 | -1 | -1 | 1 | -1 | 1 | -1 | -1 | -1 |
| RIJC024 | -1 | -1 | 1 | 1 | 1 | 1 | -1 | 1 | -1 | 1 | -1 | -1 |
| RIJC025 | -1 | 1 | 1 | -1 | -1 | -1 | -1 | -1 | 1 | -1 | 1 | 1 |
| RIJC026 | -1 | -1 | -1 | -1 | -1 | -1 | 1 | -1 | -1 | -1 | -1 | -1 |
| RIJC027 | 1 | -1 | 1 | 1 | 1 | 1 | -1 | 1 | -1 | 1 | -1 | -1 |
| RIJC029 | 1 | 1 | 1 | 1 | -1 | -1 | 1 | 1 | -1 | 1 | -1 | -1 |
| RIJC030 | -1 | -1 | -1 | -1 | -1 | -1 | -1 | -1 | 1 | -1 | 1 | 1 |
| RIJC031 | -1 | -1 | -1 | -1 | 1 | 1 | 1 | -1 | -1 | 1 | 1 | 1 |
| RIJC032 | -1 | -1 | -1 | -1 | 1 | 1 | -1 | -1 | 1 | -1 | -1 | -1 |
| RIJC045 | 1 | 1 | 1 | 1 | -1 | -1 | -1 | 1 | 1 | -1 | -1 | -1 |
| RIJC046 | 1 | -1 | -1 | -1 | 1 | 1 | 1 | 1 | -1 | 1 | -1 | 1 |
| RIJC047 | 1 | 1 | 1 | 1 | 1 | 1 | 1 | 1 | -1 | -1 | -1 | -1 |
| RIJC048 | -1 | -1 | -1 | -1 | 1 | 1 | 1 | -1 | -1 | 1 | -1 | -1 |
| RIJC049 | 1 | 1 | -1 | -1 | 1 | 1 | -1 | -1 | 1 | 1 | -1 | -1 |
| RIJC059 | -1 | -1 | 1 | 1 | 1 | 1 | -1 | -1 | -1 | 1 | 1 | 1 |
| RIJC061 | -1 | 1 | 1 | 1 | -1 | -1 | 1 | -1 | 1 | 1 | 1 | 1 |
| RIJC062 | 1 | 1 | 1 | -1 | 1 | -1 | -1 | -1 | 1 | 1 | -1 | -1 |
| RIJC064 | 1 | 1 | -1 | 1 | 1 | 1 | 1 | 1 | 1 | -1 | -1 | -1 |
| RIJC065 | 1 | -1 | -1 | 1 | 1 | 1 | 1 | 1 | -1 | -1 | -1 | -1 |
| RIJC066 | 1 | 1 | 1 | 1 | 1 | 1 | 1 | 1 | 1 | 1 | 1 | 1 |
| RIJC067 | -1 | -1 | -1 | -1 | -1 | 1 | -1 | -1 | -1 | 1 | 1 | 1 |
| RIJC069 | -1 | -1 | -1 | -1 | 1 | 1 | -1 | -1 | -1 | 1 | 1 | 1 |
| RIJC070 | 1 | -1 | -1 | -1 | -1 | -1 | -1 | -1 | -1 | 1 | 1 | 1 |
| RIJC071 | 1 | 1 | 1 | -1 | 1 | 1 | 1 | 1 | -1 | 1 | 1 | -1 |
| RIJC072 | -1 | -1 | -1 | -1 | 1 | 1 | 1 | -1 | -1 | -1 | -1 | -1 |
| RIJC073 | -1 | -1 | -1 | -1 | 1 | 1 | -1 | 1 | -1 | -1 | 1 | 1 |
| RIJC074 | 1 | 1 | 1 | 1 | 1 | 1 | 1 | -1 | 1 | -1 | 1 | -1 |
| RIJC075 | -1 | -1 | -1 | -1 | -1 | -1 | -1 | -1 | 1 | -1 | 1 | 1 |
| RIJC076 | -1 | -1 | -1 | -1 | -1 | -1 | 1 | -1 | -1 | -1 | -1 | -1 |
| RIJC078 | 1 | 1 | -1 | 1 | 1 | -1 | 1 | -1 | -1 | -1 | 1 | 1 |
| RIJC079 | -1 | 1 | 1 | 1 | 1 | 1 | -1 | -1 | -1 | 1 | 1 | 1 |
| RIJC080 | 1 | 1 | 1 | 1 | 1 | 1 | -1 | -1 | -1 | -1 | -1 | -1 |
| RIJC081 | -1 | 1 | -1 | -1 | 1 | 1 | 1 | -1 | 1 | 1 | 1 | 1 |

| | | | | | | | | | | | | |
|---|---|---|---|---|---|---|---|---|---|---|---|---|
| RIJC082 | -1 | -1 | -1 | -1 | 1 | 1 | 1 | -1 | -1 | -1 | 1 | 1 |
| RIJC129 | 1 | -1 | -1 | -1 | -1 | -1 | 1 | -1 | -1 | -1 | 1 | 1 |
| RIJC130 | 1 | 1 | -1 | -1 | -1 | -1 | 1 | 1 | -1 | 1 | -1 | 1 |
| RIJC131 | 1 | 1 | 1 | 1 | 1 | 1 | -1 | -1 | 1 | 1 | 1 | -1 |
| RIJC133 | -1 | 1 | 1 | 1 | 1 | 1 | 1 | -1 | 1 | 1 | 1 | 1 |
| RIJC135 | -1 | -1 | -1 | -1 | -1 | -1 | 1 | 1 | 1 | 1 | -1 | -1 |
| RIJC136 | -1 | -1 | -1 | -1 | 1 | 1 | -1 | -1 | -1 | 1 | -1 | -1 |
| RIJC137 | 1 | 1 | 1 | 1 | -1 | -1 | 1 | -1 | -1 | -1 | 1 | 1 |
| RIJC138 | 1 | -1 | -1 | -1 | -1 | -1 | 1 | -1 | -1 | -1 | -1 | -1 |
| RIJC139 | -1 | -1 | -1 | -1 | 1 | 1 | 1 | -1 | -1 | 1 | -1 | -1 |
| RIJC140 | 1 | 1 | 1 | 1 | -1 | -1 | 1 | 1 | -1 | -1 | 1 | 1 |
| RIJC141 | -1 | -1 | -1 | -1 | -1 | -1 | -1 | 1 | 1 | -1 | 1 | 1 |
| RIJC142 | -1 | -1 | -1 | -1 | 1 | -1 | 1 | 1 | 1 | -1 | -1 | -1 |
| RIJC144 | -1 | -1 | -1 | -1 | -1 | -1 | -1 | -1 | 1 | 1 | -1 | -1 |
| RIJC145 | -1 | -1 | -1 | 1 | 1 | 1 | -1 | 1 | 1 | -1 | -1 | -1 |
| RIJC146 | -1 | -1 | -1 | -1 | 1 | 1 | 1 | 1 | -1 | -1 | 1 | 1 |
| RIJC147 | -1 | -1 | -1 | -1 | -1 | -1 | -1 | -1 | 1 | 1 | 1 | 1 |
| RIJC148 | 1 | 1 | 1 | 1 | -1 | 1 | 1 | 1 | -1 | 1 | -1 | -1 |
| RIJC149 | -1 | -1 | -1 | -1 | 1 | 1 | 1 | -1 | -1 | -1 | 1 | 1 |
| RIJC151 | -1 | -1 | -1 | -1 | 1 | 1 | -1 | 1 | 1 | 1 | 1 | 1 |
| RIJC201 | 1 | 1 | 1 | 1 | -1 | -1 | 1 | -1 | -1 | -1 | -1 | -1 |
| RIJC202 | -1 | -1 | -1 | -1 | 1 | -1 | 1 | 1 | 1 | -1 | -1 | -1 |
| RIJC203 | -1 | 1 | -1 | -1 | -1 | -1 | 1 | -1 | 1 | 1 | 1 | 1 |
| RIJC204 | -1 | 1 | -1 | -1 | -1 | -1 | 1 | -1 | 1 | 1 | 1 | 1 |
| RIJC205 | 1 | -1 | -1 | -1 | 1 | 1 | -1 | -1 | 1 | -1 | 1 | 1 |
| RIJC206 | -1 | -1 | -1 | 1 | 1 | 1 | 1 | -1 | -1 | -1 | -1 | -1 |
| RIJC207 | -1 | -1 | -1 | -1 | 1 | 1 | -1 | -1 | -1 | 1 | 1 | 1 |
| RIJC208 | -1 | -1 | -1 | -1 | -1 | -1 | -1 | -1 | -1 | -1 | -1 | 1 |
| RIJC209 | -1 | -1 | -1 | -1 | 1 | 1 | 1 | -1 | 1 | -1 | 1 | 1 |
| RIJC210 | 1 | -1 | -1 | 1 | -1 | -1 | 1 | -1 | -1 | -1 | 1 | 1 |
| RIJC212 | 1 | -1 | 1 | 1 | 1 | 1 | 1 | -1 | -1 | -1 | 1 | 1 |
| RIJC213 | 1 | 1 | 1 | 1 | 1 | 1 | 1 | 1 | 1 | -1 | 1 | 1 |
| RIJC214 | 1 | 1 | -1 | 1 | 1 | 1 | 1 | 1 | 1 | -1 | 1 | 1 |
| RIJC216 | 1 | -1 | -1 | 1 | -1 | -1 | 1 | -1 | 1 | -1 | 1 | 1 |
| RIJC217 | 1 | 1 | 1 | -1 | -1 | -1 | -1 | -1 | -1 | -1 | 1 | -1 |
| RIJC218 | 1 | -1 | 1 | -1 | 1 | 1 | -1 | -1 | 1 | -1 | 1 | 1 |
| RIJC219 | -1 | -1 | -1 | -1 | 1 | 1 | 1 | 1 | -1 | -1 | 1 | 1 |
| RIJC220 | -1 | -1 | -1 | -1 | 1 | 1 | -1 | -1 | -1 | -1 | -1 | -1 |
| RIJC221 | 1 | -1 | 1 | -1 | 1 | -1 | 1 | 1 | 1 | 1 | -1 | 1 |
| RIJC223 | 1 | 1 | 1 | -1 | -1 | -1 | -1 | 1 | -1 | -1 | 1 | 1 |
| RIJC224 | 1 | -1 | -1 | 1 | -1 | -1 | -1 | 1 | -1 | 1 | 1 | 1 |
| RIJC225 | -1 | 1 | 1 | 1 | -1 | 1 | -1 | -1 | -1 | 1 | 1 | 1 |
| RIJC226 | -1 | -1 | 1 | 1 | 1 | 1 | -1 | 1 | 1 | 1 | 1 | 1 |
| RIJC229 | -1 | -1 | -1 | -1 | -1 | 1 | -1 | -1 | -1 | 1 | 1 | 1 |
| RIJC230 | -1 | -1 | -1 | -1 | 1 | -1 | -1 | 1 | -1 | -1 | -1 | -1 |
| RIJC231 | 1 | 1 | 1 | 1 | 1 | -1 | -1 | -1 | 1 | 1 | 1 | -1 |
| RIJC232 | -1 | -1 | -1 | -1 | -1 | -1 | 1 | 1 | 1 | -1 | -1 | -1 |
| RIJC233 | 1 | 1 | 1 | -1 | 1 | 1 | 1 | 1 | -1 | -1 | -1 | -1 |
| RIJC234 | 1 | 1 | 1 | 1 | -1 | 1 | 1 | 1 | 1 | -1 | -1 | 1 |
| RIJC235 | -1 | -1 | -1 | -1 | -1 | -1 | 1 | -1 | 1 | -1 | -1 | -1 |
| RIJC236 | -1 | -1 | -1 | -1 | -1 | -1 | 1 | 1 | -1 | 1 | -1 | -1 |
| RIJC237 | 1 | -1 | -1 | -1 | 1 | 1 | 1 | 1 | -1 | 1 | -1 | -1 |
| RIJC238 | -1 | -1 | -1 | -1 | 1 | 1 | 1 | 1 | -1 | 1 | -1 | -1 |
| RIJC242 | -1 | -1 | -1 | -1 | -1 | 1 | 1 | -1 | -1 | 1 | -1 | -1 |
| RIJC243 | -1 | -1 | -1 | -1 | 1 | 1 | 1 | -1 | -1 | -1 | -1 | -1 |
| RIJC244 | 1 | 1 | 1 | -1 | 1 | 1 | 1 | -1 | -1 | -1 | 1 | 1 |
| RIJC245 | 1 | 1 | 1 | -1 | 1 | 1 | -1 | -1 | -1 | -1 | 1 | 1 |
| RIJC247 | -1 | 1 | -1 | 1 | -1 | -1 | 1 | 1 | -1 | -1 | -1 | 1 |
| RIJC248 | 1 | 1 | 1 | 1 | -1 | -1 | 1 | 1 | -1 | -1 | 1 | 1 |
| RIJC249 | 1 | -1 | -1 | -1 | -1 | -1 | 1 | -1 | 1 | -1 | -1 | -1 |
| RIJC250 | 1 | 1 | 1 | 1 | 1 | 1 | 1 | 1 | -1 | -1 | 1 | 1 |
| RIJC251 | 1 | 1 | -1 | -1 | -1 | 1 | 1 | -1 | -1 | 1 | 1 | 1 |

| | | | | | | | | | | | | |
|---|---|---|---|---|---|---|---|---|---|---|---|---|
| RIJC252 | 1 | 1 | 1 | 1 | -1 | -1 | 1 | -1 | 1 | 1 | 1 | 1 |
| RIJC253 | 1 | 1 | 1 | 1 | -1 | -1 | -1 | -1 | 1 | 1 | 1 | 1 |
| RIJC254 | 1 | 1 | 1 | 1 | -1 | -1 | -1 | -1 | -1 | -1 | -1 | -1 |
| RIJC255 | 1 | 1 | 1 | 1 | -1 | -1 | -1 | -1 | -1 | 1 | -1 | -1 |
| RIJC256 | -1 | -1 | 1 | 1 | 1 | 1 | 1 | -1 | 1 | -1 | 1 | 1 |
| RIJC257 | 1 | 1 | 1 | 1 | -1 | 1 | 1 | -1 | -1 | -1 | -1 | -1 |
| RIJC259 | -1 | -1 | -1 | -1 | 1 | 1 | 1 | 1 | -1 | -1 | 1 | 1 |
| RIJC261 | -1 | 1 | 1 | 1 | -1 | -1 | 1 | 1 | 1 | -1 | -1 | -1 |
| RIJC262 | -1 | -1 | -1 | 1 | -1 | -1 | 1 | -1 | 1 | -1 | -1 | -1 |
| RIJC264 | -1 | -1 | -1 | -1 | 1 | 1 | 1 | -1 | 1 | 1 | 1 | 1 |
| RIJC301 | 1 | -1 | -1 | -1 | 1 | -1 | 1 | -1 | -1 | 1 | -1 | 1 |
| RIJC302 | -1 | 1 | 1 | 1 | 1 | 1 | -1 | 1 | 1 | -1 | -1 | -1 |
| RIJC303 | -1 | -1 | -1 | -1 | -1 | -1 | -1 | -1 | -1 | 1 | 1 | 1 |
| RIJC305 | -1 | -1 | -1 | -1 | -1 | -1 | 1 | 1 | 1 | -1 | -1 | -1 |
| RIJC306 | -1 | -1 | -1 | -1 | 1 | 1 | -1 | -1 | 1 | -1 | 1 | 1 |
| RIJC307 | 1 | -1 | 1 | -1 | -1 | -1 | -1 | 1 | -1 | -1 | -1 | -1 |
| RIJC309 | -1 | -1 | -1 | -1 | -1 | -1 | 1 | 1 | 1 | -1 | -1 | -1 |
| RIJC310 | 1 | 1 | 1 | 1 | 1 | 1 | -1 | 1 | -1 | -1 | -1 | -1 |
| RIJC311 | 1 | 1 | 1 | 1 | -1 | -1 | 1 | -1 | 1 | -1 | 1 | -1 |
| RIJC312 | -1 | -1 | -1 | -1 | 1 | 1 | 1 | -1 | -1 | 1 | 1 | 1 |
| RIJC313 | 1 | 1 | 1 | 1 | -1 | -1 | -1 | -1 | 1 | -1 | 1 | 1 |
| RIJC314 | -1 | 1 | 1 | 1 | -1 | -1 | 1 | -1 | -1 | 1 | 1 | 1 |
| RIJC316 | 1 | 1 | -1 | -1 | -1 | -1 | -1 | -1 | -1 | -1 | -1 | 1 |
| RIJC317 | 1 | -1 | -1 | -1 | -1 | -1 | 1 | 1 | -1 | 1 | 1 | 1 |
| RIJC318 | -1 | -1 | 1 | 1 | -1 | -1 | 1 | 1 | 1 | -1 | -1 | -1 |
| RIJC319 | -1 | 1 | -1 | 1 | -1 | -1 | -1 | 1 | -1 | 1 | -1 | -1 |
| RIJC320 | -1 | -1 | 1 | 1 | -1 | -1 | -1 | 1 | -1 | -1 | 1 | 1 |
| RIJC321 | -1 | -1 | -1 | -1 | -1 | -1 | 1 | -1 | 1 | 1 | -1 | -1 |
| RIJC322 | 1 | 1 | 1 | 1 | -1 | 1 | 1 | -1 | 1 | 1 | 1 | 1 |
| RIJC325 | -1 | -1 | -1 | -1 | 1 | 1 | 1 | 1 | 1 | 1 | 1 | 1 |
| RIJC326 | 1 | 1 | -1 | -1 | 1 | 1 | 1 | -1 | 1 | -1 | -1 | -1 |
| RIJC327 | 1 | 1 | 1 | 1 | -1 | -1 | 1 | -1 | 1 | -1 | 1 | -1 |
| RIJC328 | 1 | 1 | 1 | 1 | -1 | -1 | 1 | -1 | 1 | -1 | -1 | -1 |
| RIJC330 | -1 | -1 | -1 | -1 | -1 | -1 | -1 | -1 | -1 | -1 | 1 | -1 |
| RIJC332 | -1 | 1 | 1 | -1 | 1 | 1 | -1 | 1 | -1 | 1 | -1 | 1 |
| RIJC334 | -1 | -1 | -1 | -1 | -1 | -1 | 1 | -1 | 1 | -1 | 1 | -1 |
| RIJC335 | 1 | 1 | 1 | 1 | -1 | -1 | -1 | -1 | 1 | -1 | 1 | 1 |
| RIJC337 | 1 | -1 | -1 | -1 | -1 | 1 | -1 | -1 | 1 | 1 | -1 | 1 |
| RIJC339 | -1 | -1 | -1 | -1 | -1 | -1 | 1 | -1 | 1 | -1 | 1 | 1 |
| RIJC340 | -1 | 1 | 1 | 1 | 1 | 1 | 1 | 1 | -1 | 1 | 1 | 1 |
| RIJC341 | 1 | 1 | 1 | 1 | 1 | -1 | 1 | 1 | 1 | 1 | -1 | -1 |
| RIJC342 | -1 | 1 | -1 | -1 | -1 | 1 | 1 | -1 | 1 | -1 | -1 | -1 |
| RIJC343 | 1 | -1 | -1 | -1 | 1 | 1 | 1 | 1 | 1 | 1 | 1 | 1 |
| RIJC344 | -1 | -1 | -1 | -1 | 1 | -1 | 1 | -1 | -1 | 1 | 1 | 1 |
| RIJC346 | 1 | -1 | -1 | -1 | -1 | -1 | -1 | 1 | 1 | -1 | 1 | 1 |
| RIJC347 | -1 | -1 | -1 | -1 | -1 | -1 | 1 | 1 | 1 | 1 | -1 | 1 |
| RIJC348 | 1 | 1 | 1 | 1 | -1 | -1 | 1 | -1 | -1 | -1 | -1 | -1 |
| RIJC349 | -1 | -1 | -1 | -1 | 1 | -1 | 1 | 1 | -1 | -1 | 1 | 1 |
| RIJC350 | -1 | -1 | -1 | -1 | -1 | -1 | -1 | -1 | 1 | -1 | 1 | 1 |
| RIJC351 | -1 | -1 | -1 | -1 | -1 | -1 | 1 | -1 | 1 | -1 | -1 | -1 |
| RIJC352 | -1 | -1 | 1 | 1 | 1 | 1 | -1 | 1 | -1 | 1 | 1 | -1 |
| RIJC353 | -1 | 1 | 1 | 1 | 1 | 1 | -1 | 1 | -1 | 1 | -1 | -1 |
| RIJC354 | 1 | 1 | 1 | 1 | 1 | 1 | 1 | 1 | -1 | -1 | -1 | -1 |
| RIJC355 | 1 | -1 | 1 | 1 | 1 | 1 | -1 | 1 | 1 | -1 | -1 | -1 |
| RIJC356 | -1 | -1 | -1 | -1 | 1 | 1 | -1 | -1 | 1 | -1 | 1 | 1 |
| RIJC357 | -1 | -1 | -1 | -1 | 1 | 1 | 1 | 1 | -1 | -1 | -1 | -1 |
| RIJC358 | -1 | -1 | 1 | 1 | -1 | -1 | 1 | 1 | -1 | -1 | -1 | -1 |
| RIJC360 | 1 | -1 | 1 | -1 | -1 | -1 | 1 | -1 | 1 | -1 | -1 | -1 |
| RIJC361 | -1 | -1 | 1 | 1 | -1 | -1 | -1 | -1 | -1 | -1 | -1 | -1 |
| RIJC362 | -1 | -1 | -1 | -1 | 1 | 1 | 1 | -1 | 1 | 1 | 1 | -1 |
| RIJC363 | 1 | 1 | 1 | 1 | 1 | 1 | 1 | -1 | -1 | 1 | 1 | 1 |
| RIJC364 | 1 | 1 | 1 | 1 | -1 | 1 | -1 | 1 | -1 | 1 | 1 | 1 |

| | | | | | | | | | | | | |
|---|---|---|---|---|---|---|---|---|---|---|---|---|
| RIJC366 | 1 | 1 | -1 | -1 | 1 | 1 | 1 | -1 | 1 | -1 | -1 | -1 |
| RIJC367 | 1 | 1 | 1 | 1 | 1 | 1 | -1 | -1 | 1 | 1 | 1 | 1 |
| RIJC368 | 1 | 1 | 1 | 1 | -1 | -1 | -1 | 1 | -1 | 1 | -1 | -1 |
| RIJC369 | 1 | 1 | 1 | 1 | -1 | -1 | -1 | 1 | 1 | -1 | 1 | 1 |
| RIJC370 | -1 | -1 | 1 | 1 | 1 | -1 | -1 | -1 | 1 | -1 | -1 | -1 |
| RIJC371 | -1 | -1 | -1 | -1 | -1 | 1 | -1 | 1 | 1 | -1 | -1 | -1 |
| RIJC372 | 1 | 1 | 1 | 1 | -1 | -1 | -1 | 1 | -1 | 1 | 1 | -1 |
| RIJC373 | -1 | -1 | -1 | -1 | -1 | -1 | -1 | -1 | -1 | -1 | 1 | 1 |
| RIJC374 | 1 | 1 | -1 | -1 | -1 | -1 | 1 | -1 | -1 | 1 | -1 | -1 |
| RIJC375 | -1 | 1 | -1 | 1 | -1 | -1 | 1 | -1 | 1 | 1 | 1 | 1 |

**Table S2.** Estimated terms in the dynamic QTL effect module showing the estimated parameter values with confidence intervals and p-value for the rate of progress from planting to flowering.

| Terms [a] | Estimated | 2.5% | 97.5% | p-value |
|---|---|---|---|---|
| Intercept | $2.35148 \times 10^{-2}$ | $2.33418 \times 10^{-2}$ | $2.36879 \times 10^{-2}$ | $9.01155 \times 10^{-232}$ |
| $Tmax_{s,g}$ | $5.72311 \times 10^{-4}$ | $5.11997 \times 10^{-4}$ | $6.32626 \times 10^{-4}$ | $4.37596 \times 10^{-62}$ |
| $Tmin_{s,g}$ | $5.29789 \times 10^{-4}$ | $4.87553 \times 10^{-4}$ | $5.72025 \times 10^{-4}$ | $1.32239 \times 10^{-94}$ |
| $DayL_{s,g}$ | $-1.56357 \times 10^{-3}$ | $-1.65041 \times 10^{-3}$ | $-1.47673 \times 10^{-3}$ | $1.53907 \times 10^{-152}$ |
| $Srad_{s,g}$ | $-8.50211 \times 10^{-5}$ | $-1.20053 \times 10^{-4}$ | $-4.99896 \times 10^{-5}$ | $2.42422 \times 10^{-6}$ |
| $QTL1_{s,g}$ | $9.41278 \times 10^{-4}$ | $7.67693 \times 10^{-4}$ | $1.11486 \times 10^{-3}$ | $1.14674 \times 10^{-20}$ |
| $QTL2_{s,g}$ | $1.24887 \times 10^{-3}$ | $1.05220 \times 10^{-3}$ | $1.44554 \times 10^{-3}$ | $6.18612 \times 10^{-26}$ |
| $QTL3_{s,g}$ | $-6.08364 \times 10^{-4}$ | $-8.29898 \times 10^{-}4$ | $-3.86831 \times 10^{-4}$ | $2.26795 \times 10^{-7}$ |
| $QTL4_{s,g}$ | $2.36803 \times 10^{-4}$ | $4.20996 \times 10^{-5}$ | $4.31506 \times 10^{-4}$ | $1.81926 \times 10^{-2}$ |
| $QTL5_{s,g}$ | $5.67194 \times 10^{-6}$ | $-1.86555 \times 10^{-4}$ | $1.97899 \times 10^{-4}$ | $9.53950 \times 10^{-1}$ |
| $QTL6_{s,g}$ | $5.27617 \times 10^{-4}$ | $3.36592 \times 10^{-4}$ | $7.18642 \times 10^{-4}$ | $2.07379 \times 10^{-7}$ |
| $QTL7_{s,g}$ | $-4.11459 \times 10^{-4}$ | $-5.56487 \times 10^{-4}$ | $-2.66430 \times 10^{-4}$ | $9.99240 \times 10^{-8}$ |
| $QTL8_{s,g}$ | $-2.11983 \times 10^{-4}$ | $-3.60161 \times 10^{-4}$ | $-6.38048 \times 10^{-5}$ | $5.61825 \times 10^{-3}$ |
| $QTL9_{s,g}$ | $-4.42610 \times 10^{-4}$ | $-5.83805 \times 10^{-4}$ | $-3.01415 \times 10^{-4}$ | $5.28426 \times 10^{-9}$ |
| $QTL10_{s,g}$ | $-2.50138 \times 10^{-4}$ | $-3.95428 \times 10^{-4}$ | $-1.04847 \times 10^{-4}$ | $9.14723 \times 10^{-4}$ |
| $QTL11_{s,g}$ | $3.43389 \times 10^{-4}$ | $1.22771 \times 10^{-4}$ | $5.64006 \times 10^{-4}$ | $2.63477 \times 10^{-3}$ |
| $QTL12_{s,g}$ | $-1.53677 \times 10^{-4}$ | $-3.72048 \times 10^{-4}$ | $6.46939 \times 10^{-5}$ | $1.69555 \times 10^{-1}$ |
| $QTL1_{s,g} \times QTL2_{s,g}$ | $3.01579 \times 10^{-4}$ | $1.27364 \times 10^{-4}$ | $4.75794 \times 10^{-4}$ | $8.55786 \times 10^{-4}$ |
| $DayL_{s,g} \times QTL3_{s,g}$ | $-7.66441 \times 10^{-4}$ | $-8.44228 \times 10^{-4}$ | $-6.88654 \times 10^{-4}$ | $4.92482 \times 10^{-66}$ |
| $DayL_{s,g} \times QTL7_{s,g}$ | $-1.62459 \times 10^{-4}$ | $-2.21472 \times 10^{-4}$ | $-1.03446 \times 10^{-4}$ | $9.57607 \times 10^{-8}$ |
| $DayL_{s,g} \times QTL12_{s,g}$ | $-1.45956 \times 10^{-4}$ | $-2.11416 \times 10^{-4}$ | $-8.04965 \times 10^{-5}$ | $1.44756 \times 10^{-5}$ |
| $Tmin_{s,g} \times QTL2_{s,g}$ | $-2.40896 \times 10^{-5}$ | $-5.95337 \times 10^{-5}$ | $1.13545 \times 10^{-5}$ | $1.83307 \times 10^{-1}$ |
| $Tmin_{s,g} \times QTL3_{s,g}$ | $-8.59042 \times 10^{-5}$ | $-1.28746 \times 10^{-4}$ | $-4.30624 \times 10^{-5}$ | $9.42280 \times 10^{-5}$ |
| $Tmax_{s,g} \times QTL5_{s,g}$ | $8.54093 \times 10^{-5}$ | $3.87125 \times 10^{-5}$ | $1.32106 \times 10^{-4}$ | $3.62986 \times 10^{-4}$ |
| $Srad_{s,g} \times QTL12_{s,g}$ | $-3.06273 \times 10^{-5}$ | $-6.04701 \times 10^{-5}$ | $-7.84445 \times 10^{-7}$ | $4.46841 \times 10^{-2}$ |
| Fixed effects variance | $1.80182 \times 10^{-5}$ | | | |
| Random effects variance | $6.34775 \times 10^{-7}$ | | | |
| Residual variance | $1.30567 \times 10^{-6}$ | | | |

[a] Predictor variable are: Intercept represents the overall value of daily development rate, maximum and minimum temperature (Tmax and Tmin, °C), day length (DayL, hours) and solar radiation (Srad, MJ $m^{-2}$ $d^{-1}$). The rate of progress to flowering rate is (1/duration from plating to flowering in days) for the $g^{th}$ genotype for the $s^{th}$ site; QTL1 to QTL12 allele effects are represented as +1 for Calima alleles and -1 for Jamapa alleles.

***Figure S1*** Parameter estimation process for predicting first flowering across all sites using the Dynamic Mixed Linear Module (DMLM

```
## cleaning memory and loading R libraries
rm(list=ls())
library("readr")
library(stats)
library(lmerTest)
library(tidyverse)

### Setup your directory location
setwd("C:\path\to\my\directory")

##Environmental data
we <- read.table("C:\path\to\environment\data",header=T,sep=',')

# Example of table structure to read the environmental data.
# (data.frame(SITE = character(),
#             Srad = numeric(),
#             Tmax = numeric(),
#             Tmin = numeric(),
#             DayL = numeric()))

##Recombinant inbred lines (see Table S1)
r1 <- read_csv("C:\path\to\QTL\data")


## Part 1 - Parameter estimation

## First of all, provide the observed days to first flowering for each RIL
## and store it as a new column (r1$R1). Then extract the mean of
## environmental data from sowing ## to first flower for each RIL and store
## it as a new column for r1$Sradm, r1$Tminm, r1$Tmaxm and r1$DayLm.


## Mean of environmental data from sowing to first flower across all
## genotypes, sites, years.
#sradMean <- mean(r1$Sradm)
#dayMean  <- mean(r1$DayLm)
#tminMean <- mean(r1$Tminm)
#tmaxMean <- mean(r1$Tmaxm)


##Centering continuous variable
r1$Srad_c <-r1$Sradm - sradMean
r1$Day_c  <-r1$DayLm - dayMean
r1$Tmin_c <-r1$Tminm - tminMean
r1$Tmax_c <-r1$Tmaxm - tmaxMean

### 1/days to first flowering
r1$R1rate <- 1/r1$R1

##Dynamic mixed-effects linear model for first flowering rate
modelr1rate = lmer((R1rate)~
                     Tmax_c+
                     Tmin_c+
```

```
                     Day_c+
                     Srad_c+
                     QTL1+QTL2+QTL3+QTL4+QTL5+QTL6+
                     QTL7+QTL8+QTL9+QTL10+QTL11+QTL12+
                     QTL1*QTL2+
                     Day_c*QTL3+
                     Day_c*QTL7+
                     Day_c*QTL12+
                     Tmin_c*QTL2+
                     Tmin_c*QTL3+
                     Tmax_c*QTL5+
                     Srad_c*QTL12+
                     (1|RIL),data=r1)
summary(modelr1rate)

## Part 2 - Estimating daily flowering rate for each RIL at each SITE
predictionlist <- list()
predictionlist[[1]] =
rbind(predictionlist, c("SITE","RIL",'Observed','Simulated'))
rowcount = 1
stepper <-1

## Change the site based on your data
si <-  c("ND","FL","PR","PA","PO")
for (s in si){

  r2<-subset(r1,r1$SITE==s)  # subset QTL data file
  we1 <- subset(we,SITE==s)  # subset weather data file

  for (i in 1:nrow(r2)) {
    SITE  <- c(as.character(r2$SITE[i]))
    rsite <- SITE
    RIL   <- c(as.character(r2$RIL[i]))
    Observed <- c(r2$R1[i])
    QTL1  <- c(r2$QTL1[i])
    QTL2  <- c(r2$QTL2[i])
    QTL3  <- c(r2$QTL3[i])
    QTL4  <- c(r2$QTL4[i])
    QTL5  <- c(r2$QTL5[i])
    QTL6  <- c(r2$QTL6[i])
    QTL7  <- c(r2$QTL7[i])
    QTL8  <- c(r2$QTL8[i])
    QTL9  <- c(r2$QTL9[i])
    QTL10 <- c(r2$QTL10[i])
    QTL11 <- c(r2$QTL11[i])
    QTL12 <- c(r2$QTL12[i])

    CounterR1 = 0
    DayCount = 0


    for (j in 1:nrow(we1)){
      DayCount <- we1$DAP[j]

      # Adjusting weather variables
      Srad_c <- c(we1$Srad[j])-sradMean
      Day_c <- c(we1$DayL[j])-dayMean
```

```
      Tmax_c <- c(we1$Tmax[j])-tmaxMean
      Tmin_c <- c(we1$Tmin[j])-tminMean
      r1day <- as.data.frame(SITE,RIL,QTL1,QTL2,QTL3,QTL4,QTL5,
                             QTL6,QTL7,QTL8,QTL9,QTL10,QTL11,QTL12,
                             Srad_c,Tmin_c,Tmax_c,Day_c)

      ## Estimating daily flowering rate gain based on "modelr1rate" model
      flowerNow <- predict(modelr1rate,re.form=NA,newdata=r1day)

      ## counting cumulative flowering rate
      CounterR1 = CounterR1 + flowerNow[[1]]
      dailyRateR1 = flowerNow[[1]]
      rowcount = rowcount + 1

      ## exiting loop when rate reaches 1.00
      if (CounterR1[[1]] >= 1.00){
        Simulated <- DayCount
        predictionlist[[rowcount]] <- c(SITE,RIL,Observed,Simulated)
        break}
    }

    stepper = stepper +1
    print(c(SITE,stepper))
  }
}

# converting list to matrix
predictionRate <- as.matrix(do.call("rbind", predictionlist))
colnames(predictionRate)<-predictionRate[1,]  # fixing header row names
predictionRate<-predictionRate[-1,] # removing old header row

##View daily rate prediction for QTL allele combination
View(predictionRate)

#END
```

***Figure S1*** Computer code for the Dynamic Piecewise Linear Module (DPLM) coupled with the CSM-CROPGRO-Drybean model

```
!=======================================================================
!  Dynamic piecewise linear module, Program,
!=======================================================================
!-----------------------------------------------------------------------

      SUBROUTINE DPLM(CONTROL, ISWITCH,                    &  !Control
          WEATHER, YRPLT,                                  &  !Input
          NR1G, SumFRD)                                       !Output

!-----------------------------------------------------------------------
      USE ModuleDefs

      IMPLICIT none
      SAVE
!-----------------------------------------------------------------------
      CHARACTER*6   GENID
      CHARACTER*30 FILEIO

      LOGICAL FRSTFL

      INTEGER RUN, DYNAMIC, DAS, YRDOY, YR, DOY, YRPLT
      INTEGER DAP, TIMDIF, FDOY, NR1G

      REAL DAYL, SRAD, TMAX, TMIN
      REAL SumFRD, FR
      REAL FRMAX, DLm, Sradm, Tmaxm, Tminm
      REAL, DIMENSION(70) :: QTL

      TYPE (ControlType) CONTROL
      TYPE (SwitchType) ISWITCH
      TYPE (WeatherType) WEATHER

      DYNAMIC = CONTROL % DYNAMIC
      FILEIO  = CONTROL % FILEIO
      RUN     = CONTROL % RUN
      DAS     = CONTROL % DAS
      YRDOY   = CONTROL % YRDOY


!***********************************************************************
!***********************************************************************
!     Run Initialization - Called once per simulation
!***********************************************************************
      IF (DYNAMIC .EQ. RUNINIT) THEN
!-----------------------------------------------------------------------
!       Read Genetic input data
!-----------------------------------------------------------------------
        CALL IPGENE(FILEIO, TF, GENID)

!***********************************************************************
!***********************************************************************
!     Seasonal initialization - run once per season
!***********************************************************************
```

```
      ELSEIF (DYNAMIC .EQ. SEASINIT) THEN
!-----------------------------------------------------------------------
!     Set sowing/start day of year for flowering model to start
!     Initialize progress toward flowering, SumFRD, & Day of First Flower
!-----------------------------------------------------------------------
        SumFRD = 0.0
        DAYL    = 0.0
        SRAD    = 0.0
        TMAX    = 0.0
        TMIN    = 0.0
        FDOY    = 0
        NR1G    = 0
        FRSTFL  = .FALSE.
        GENID   = ''
!-----------------------------------------------------------------------
!     Averages across 5 environments in datasets used to estimate model
!     mean values of environmental variables.
!-----------------------------------------------------------------------
        DLm    = 12.722559638877
        Sradm  = 18.2718980213904
        Tmaxm  = 27.4529030160428
        Tminm  = 16.1181873475936

!-----------------------------------------------------------------------
!     Limit maximum rate for a genotype based on QTLs, (FRMAX)
!-----------------------------------------------------------------------
        FRMAX  = 0.02798564527544710                &
               + 0.00107126855594358 * QTL(1)       &
               + 0.00124937987119220 * QTL(2)       &
               - 0.00035350501343249 * QTL(3)       &
               + 0.00039945509894467 * QTL(4)       &
               + 0.00007085168560867 * QTL(5)       &
               + 0.00056306276150221 * QTL(6)       &
               - 0.00042854934463711 * QTL(7)       &
               - 0.00023509947596009 * QTL(8)       &
               - 0.00060905231969296 * QTL(9)       &
               - 0.00029702065347147 * QTL(10)      &
               + 0.00063538481240068 * QTL(11)      &
               - 0.00023169840751149 * QTL(12)

!***********************************************************************
!***********************************************************************
!     Daily Rate calculations
!***********************************************************************
      ELSE IF (DYNAMIC .EQ. RATE) THEN
!----------------------------------------------------------------------
!       Read Weather data
!----------------------------------------------------------------------
        CALL YR_DOY(YRPLT,YR,DOY)
        DAP = MAX (0, TIMDIF(YRPLT, YRDOY))

        SRAD = WEATHER%SRAD
        TMAX = WEATHER%TMAX
        TMIN = WEATHER%TMIN
        DAYL = WEATHER%DAYL
!-----------------------------------------------------------------------
!       The dynamic gene-based mixed effects linear model, Bean
```

```
!-----------------------------------------------------------------------
         FR = 0.02351482064754700                                      &
             + 0.00057231114859053 * (TMAX - Tmaxm)                    &
             + 0.00052978909146124 * (TMIN - Tminm)                    &
             - 0.00156356767055738 * (DAYL - DLm)                      &
             - 0.00008502111815422 * (SRAD - Sradm)                    &
             + 0.00094127790142489 * QTL(1)                            &
             + 0.00124887042689091 * QTL(2)                            &
             - 0.00060836426965787 * QTL(3)                            &
             + 0.00023680275595848 * QTL(4)                            &
             + 0.00000567194207821 * QTL(5)                            &
             + 0.00052761690606345 * QTL(6)                            &
             - 0.00041145860322971 * QTL(7)                            &
             - 0.00021198302954652 * QTL(8)                            &
             - 0.00044260992328382 * QTL(9)                            &
             - 0.00025013760365754 * QTL(10)                           &
             + 0.00034338877075107 * QTL(11)                           &
             - 0.00015367693845455 * QTL(12)                           &
             + 0.00030157871435221 * QTL(1) * QTL(2)                   &
             - 0.00076644059984278 * (DAYL - DLm) * QTL(3)             &
             - 0.00016245875627569 * (DAYL - DLm) * QTL(7)             &
             - 0.00014595603452997 * (DAYL - DLm) * QTL(12)            &
             - 0.00002408961229346 * (TMIN - Tminm) * QTL(2)           &
             - 0.00008590422032661 * (TMIN - Tminm) * QTL(3)           &
             + 0.00008540931051535 * (TMAX - Tmaxm) * QTL(5)           &
             - 0.00003062728932614 * (SRAD - Sradm) * QTL(12)

     ! Note that one can replace the above mixed effects linear model with
     ! any function that computes each day's rate of progress toward
     ! first flowering

!***********************************************************************
!***********************************************************************
!     Daily integration
!***********************************************************************
      ELSEIF (DYNAMIC .EQ. INTEGR) THEN

!-----------------------------------------------------------------------
!     Compute time integral of development to pass back as cumulative
!       progress toward development each day
!     In the equation for computing SumFRD, the time step is assumed
!       to be 1.0 d for this module (fixed)
!-----------------------------------------------------------------------
!     Limit rate of development to positive values;
!       initial value=0.0. When SumFRD first reaches 1.00,
!       flowering will occur
!-----------------------------------------------------------------------

        IF (FR < 1E-5) THEN
          FR = 0.0
        ENDIF

        IF (FR > FRMAX) THEN
          FR = FRMAX
        ENDIF

        SumFRD = SumFRD + FR*1.0
```

```
        IF (SumFRD >= 1.0 .AND. FDOY < 1) THEN   !First flower occurs
          FRSTFL = .TRUE.
          FDOY = DAP + DOY
          NR1G = DAS
        ENDIF

!***********************************************************************
!***********************************************************************
!     END OF DYNAMIC IF CONSTRUCT
!***********************************************************************
      ENDIF

      RETURN
    END ! DPLM

!-----------------------------------------------------------------------
!     DPLM VARIABLES LIST
!-----------------------------------------------------------------------
! DAP        Number of days after planting (d).
! DAS        Days after start of simulation (d).
! DAYL       Day length on day of simulation (from sunrise to sunset) (hr).
! DLm        Mean day length across all five sites, all genotypes.
! DOY        Current day of simulation (d).
! DYNAMIC    Module control variable.
! FDOY       Number of days after planting when the first flower occurs (d).
! FRSTFL     Flag to identify that the first flower occurs (true/false).
! GENID      Identifier for reading in the input file '.gen'.
! NR1G       Number of days after start of simulation when the first flower
!            occurs (d).
! FR         Daily rate of progress from planting to first flower appearance
!            for selected genotype and environmental factors on the current
!            environment & day.
! FRMAX      Maximum rate of progress toward first flower.
! SRAD       Solar radiation (MJ/m2-d).
! Sradm      Mean solar radiation transplanting to first flower across all
!            genotypes, sites, years.
! SumFRD     Current progress toward first flowering of FR.
! QTL(n)     Alleles at QTL(1) : QTL(n) in jth genotype.
! TIMDIF     Integer function which calculates the number of days between
!            two Julian dates (da).
! TMAX       Maximum daily temperature (Celsius).
! Tmaxm      Mean of maximum temperature from transplanting to first flower
!            across all genotypes, sites, years.
! TMIN       Minimum daily temperature (Celsius)
! Tminm      Mean of minimum temperature from transplanting to first flower
!            across all genotypes, sites, years.
! YR         Year portion of date
! YRDOY      Current day of simulation (YYYYDDD)
! YRPLT      Planting date (YYYYDDD)
```
